# Quantifying the impact of genetic mutations on enhancer dynamics

**DOI:** 10.1101/2025.10.21.683558

**Authors:** Maitreya Das, Deepro Banerjee, Ayaan Hossain, Matthew Jensen, Saie Mogre, Jiawan Sun, Joseph Mao, Adam B. Glick, Howard M. Salis, Santhosh Girirajan

**Author notes:** Correspondence: Santhosh Girirajan, 205A Life Sciences Building, Pennsylvania State University. Maitreya Das and Deepro Banerjee contributed equally to this work.

## Abstract

Transcriptional regulation is mediated by enhancers, yet how genetic perturbations alter enhancer activity and gene expression remains poorly understood. We developed UDI-UMI-STARR-seq, which integrates dual indexes and unique molecular identifiers, and combined it with RNA-seq to profile the effects of perturbations on enhancer activity and target gene expression. We applied this approach to a library of 253,632 fragments representing 46,142 cell type–specific candidate enhancers and assessed the impact of CRISPR/Cas9-mediated deletion of six transcription factors (or TFs; ATF2, CTCF, FOXA1, LEF1, TCF7L2, and SCRT1) with diverse regulatory roles. Across knockout lines, we identified responsive enhancers that were either repressed or induced, often through motifs such as the p53 family of TFs. Enhancer–gene mapping revealed TF-specific programs, including repression of Wnt/p53 targets with ATF2 or LEF1 loss, downregulation of the FIRRE locus with CTCF loss, and compensatory upregulation of RNA polymerase II regulators following FOXA1 depletion. A deep learning model trained on enhancer sequences recapitulated core principles of enhancer grammar, including cooperative motif syntax and the influence of flanking sequence context. Applying this framework to the neurodevelopmental disorder-associated 16p12.1 deletion identified responsive enhancers linked to genes involved in axon guidance, synaptic plasticity, and translational control, providing a scalable readout of enhancer dynamics generalizable to genetic mutations.

## INTRODUCTION

Cellular function relies on precise, context-specific expression of genes, orchestrated by transcription factors, enhancers, and promoters, which together form the core of gene regulatory network^1–3^. Enhancers are short DNA sequences that act as binding sites for TFs, typically through 12–20 bp motifs, and physically loop over to target promoters and initiate transcription^4^. Genetic or cellular perturbation can disrupt the gene regulatory network, leading to altered gene expression and downstream functional consequences (**Fig. 1A**). For example, TP53-responsive enhancer reporter screens showed that activating or blocking a TF modulate enhancer activity and thereby influence transcription^5^. Similarly, genetic variant-level reporter assays revealed allele-specific effects on enhancer activity linked to changes in target gene expression^6^. External stimuli such as hormones or drugs can also modulate enhancer activity, reshaping downstream transcriptional responses^7,8^. Despite these insights, most studies to date emphasize on transcriptomic outcomes of perturbations or stimuli, focusing on the effect rather than the causal regulatory dynamics that produce it. Bridging this gap is essential for elucidating the principles of gene regulation, understanding its mis-regulation in disease, and mapping the broader connectivity of gene regulatory networks^9–11^.

**Figure 1:**
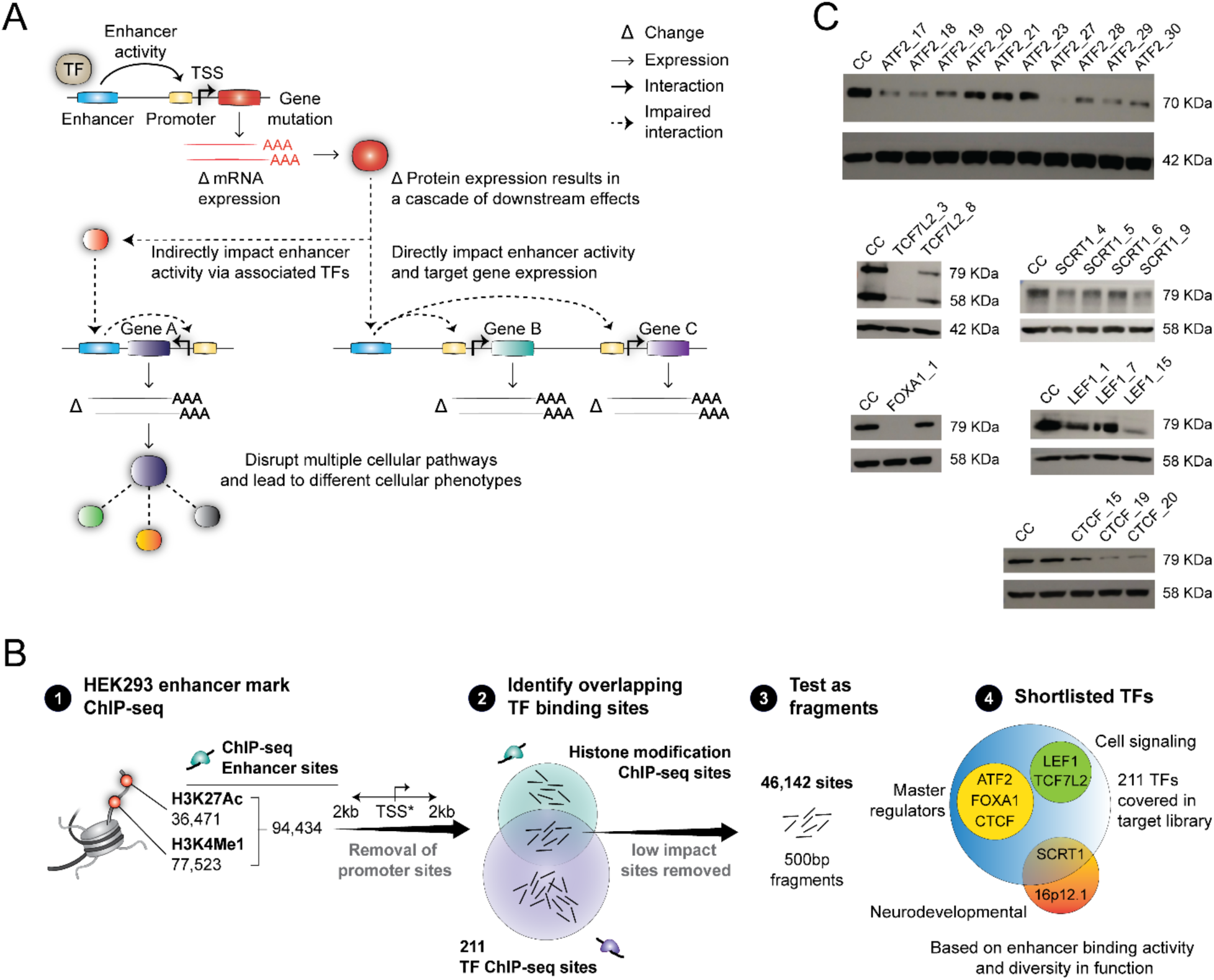
A framework to assess the effect of mutations on enhancer activity. **(A)** Schematic demonstrating the impact of genomic mutations on enhancer activity and its cascade of downstream effects. A mutation on a transcriptions factor (TF) gene can alter mRNA and protein levels, affecting TF binding to target enhancers and directly changing downstream gene expression and pathways. Alternatively, it can disrupt co-factor complexes, indirectly affecting distal regulatory element activity and target gene expression. **(B)** Flowchart showing the steps for selecting candidate enhancers from HEK293 ChIP-seq sites and shortlisting six transcription factors (TFs) from the 211 in the target library, based on enhancer binding properties, functional diversity, and ChIP-seq site overlap. **(C)** Western blots confirming CRISPR/Cas9 mediated deletion of the selected six TFs.

Assaying enhancer activity has been particularly challenging because their function is context dependent. The same enhancer can regulate different genes in distinct cellular environments and can also act cooperatively with other enhancers^12–14^. Enhancer activity is further influenced by chromatin accessibility, TF availability, compensatory TF mechanisms, and cell-type specificity, making it difficult to establish universal rules that govern enhancer responses to cellular changes. To address these challenges, high-throughput methods such as ATAC-seq, ChIP-seq, CAGE, Hi-C, MPRAs, and Perturb-seq studies have been employed to map enhancers, identify their target genes, and quantify activity across cell types^15,16^. These studies have provided important insights into enhancer-TF interactions, stimulus-dependent enhancer activation, and functional consequences of enhancer mutations^17,18^. However, methodological limitations have hindered efforts to systematically predict enhancer responses to genomic changes.

Here, we present a framework integrating UDI-UMI-STARR-seq, an adaptation of Self-Transcribing Active Regulatory Region Sequencing (STARR-seq) assay^19^, with RNA-seq to investigate the impact of gene disruption on enhancer activity. UDI-UMI-STARR-seq allows direct quantification of enhancer activity by accurately measuring self-transcribed mRNA obtained from enhancer fragments. Because TFs are the primary sequence-specific regulators of enhancer activity, we hypothesized that performing UDI-UMI-STARR-seq in TF-depleted cells would reveal high-effect and broad shifts in enhancer activity compared to disruption of other gene classes. Furthermore, TFs exert diverse regulatory roles, including opening chromatin, recruiting transcriptional co-factors, and mediating enhancer-promoter communication. Consequently, TF disruption would induce regulatory changes that are highly informative for understanding the impact of different types of genomic mutations.

We tested the impact of deleting six TFs on the activity of 253,632 candidate enhancers spanning 46,142 cell-type-specific ChIP-seq sites. To further interpret these effects, we leveraged deep learning methods to learn context-specific enhancer grammar, capturing how enhancer activity is influenced by a single motif or specific motif combinations, demonstrating how sequence features drive regulatory outcomes. Finally, we applied our framework to a neurodevelopmental disorder-associated 16p12.1 deletion, identifying responsive enhancers and their downstream targets to reconstruct a deletion-specific regulatory network. Our study provides a scalable strategy to dissect enhancer function, predict mutation-induced regulatory changes, and refine models of gene regulation in health and disease.

## RESULTS

We curated ChIP-seq datasets from ENCODE and designed a targeted library of 46,142 candidate enhancer sites (approximately 33 Mbp) that were bound by 211 TFs or having H3K4Me1 and H3K27Ac histone modifications **(Fig. 1B; Methods)**. Using CRISPR/Cas9 dual gRNA strategy^20^, we generated six knockout (KO) lines and confirmed protein depletion for transcriptional regulators *FOXA1*, *ATF2* and *CTCF*, signaling molecules *LEF1*, *TCF7L2* and *SCRT1*, compared to a non-KO control (CC) in HEK293T cells **(Fig. 1B and C; Supplementary Table 1; Supplementary Information).** These transcription factors are associated with a wide range of functions, including the ability to unmask closed chromatin and promote gene transcription, modulate the activation and repression of Wnt signaling genes, and prevent gene activation by “blocking” enhancer activity (**Supplementary Table 1)**. We assessed the impact of TF depletion on enhancer activity and gene expression by using an adaptation of STARR-seq, called UDI-UMI-STARR-seq and RNA-seq, respectively, for each KO line and CC **(Fig. 2A; Methods; Supplementary Information)**. UDI-UMI-STARR-seq was modified from CapSTARR-seq^6,21^ and UMI-STARR-seq protocols^22^, and included the target library captured from sheared whole genome DNA using custom designed hybridization and capture probes. All input and output sequencing libraries were added with Unique Molecular Identifiers (UMIs) and Illumina Unique Dual Indexes (UDIs) to enable detection of PCR duplicates and index-hopping events, respectively^23,24^ **(Fig. 2B; Methods; Supplementary Information)**. Prior to peak calling, we developed a package called STARRDUST to process, demultiplex, and deduplicate raw sequence reads **(Methods)**. The inter-replicate read counts and fold change between input DNA and output mRNA transcripts across all libraries showed strong correlations (Pearson coefficient 0.80–0.99), indicating high data quality **(Fig. 2C)**.

**Figure 2:**
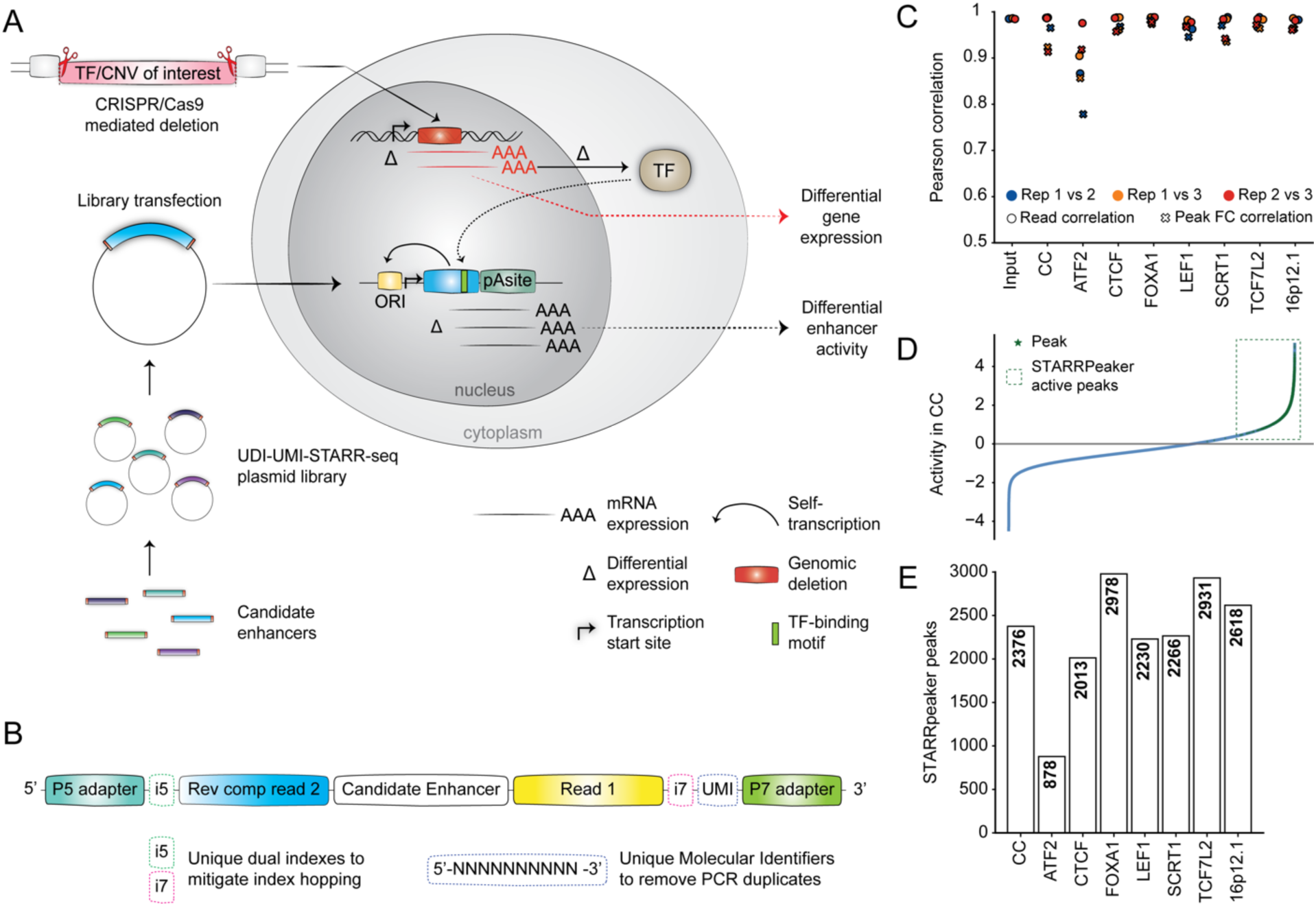
Using UDI-UMI-STARR-seq to detect differential enhancer activity. **(A)** Schematic demonstrating strategy used to assess the impact of genetic mutations using UDI-UMI-STARR-seq and RNA-seq. A plasmid library comprising >46,000 candidate enhancers was transfected into HEK293T cells in wild-type and CRISPR/Cas9-mediated knockout (KO; transcription factor and copy number variant deletion) states. Enhancer activity and gene expression was assessed using UDI-UMI-STARR-seq and RNA-Seq respectively. Each KO line was compared to wild type to quantify the effect of genetic mutations. **(B)** Modified sequence architecture used for all STARR-seq libraries. Illumina UDI and UMIs were added to all STARR-seq libraries to ensure detection of index hopping events and PCR duplicates. Both library types (input and output) carried identical architectures to ensure accurate enhancer activity assessment. **(C)** Pearson correlation coefficients across technical replicates (N=3) for the number of reads aligned to each candidate enhancer fragment and the fold change (output reads/input reads) of STARRPeaker peaks for CC and each KO library. **(D)** Observed fold change across 253,632 overlapping fragments in CC. Active enhancer peaks and fragments overlapping with STARRPeaker called peaks are highlighted as a dotted box. **(E)** Bar plot showing the number of active peaks called using STARRPeaker across CC and KO libraries.

To address target region size-induced anomalies reported in STARR-seq studies^25^ and to achieve uniform assessment across libraries, we divided the candidate library into 253,632 overlapping fragments of 500-bp size. We used STARRPeaker^26^ to define active enhancers and quantified the activity of each fragment by comparing normalized read depth of output libraries to the input **(Fig. 2D; Methods; Supplementary Information)**. As expected, active enhancers showed higher STARR-seq activity than control exonic fragments (*P*<1×10^−100^) and overlapped with open chromatin regions identified by DNase-seq in HEK293T cells, consistent with findings in prior studies^27^ **(Supplementary Fig. 1A and B)**. We identified 2,376 active fragments for CC, comprising 0.9% of all library fragments, and varying number of active fragments across all KO lines **(Fig. 2E)**. While we observed decreased number of active fragments compared to CC with the knockout of SCRT1, CTCF, LEF1, and ATF2 lines, the number of active fragments increased with the knockout of FOXA1 and TCF7L2 **(Fig 2E; Methods; Supplementary Information)**. This observation suggests that the intrinsic activity of potential enhancer fragments may change based on specific genomic perturbations. Therefore, in addition to WT cells, measuring enhancer response to genetic changes can be an additional metric to understand the properties of enhancers.

### Quantifying differential enhancer activity

We quantified the effect of each TF deletion by calculating differential enhancer activity, defined as the change in activity of each fragment in the KO lines relative to CC. We defined fragments with altered activity as ‘responsive’, whereas those unaffected by TF deletion as non-responsive, which were either consistently active or consistently inactive in all KO lines and CC **(Supplementary Data 1)**. For instance, using motif enrichment analysis, we found that the consistently active fragments were enriched for binding motifs of CHOP, ATF, JUN, FOS and the ETS family of TF that are related to cell growth, metabolism, differentiation, and apoptosis functions, as well as motifs for YY1, that is associated with neurodevelopmental processes^28^ **(Supplementary Data 2)**. In contrast, fragments that were consistently inactive contained motifs for BATF3, DLX3, MAF and HINFP, suggesting cell type-specific effects that are not apparent in HEK293T cells^29–32^. The number of responsive fragments varied across the KO lines. For instance, we observed higher numbers of responsive fragments for FOXA1 (14%, 35,245/253,632) and ATF2 KO lines (8%, 20,987/253,632), reflecting their global roles as a pioneering factor in chromatin modeling and as a core member of the AP1 family of regulators in the cell ^33^, respectively. In contrast, only 1.7% (4,288/253,632) of the fragments in the SCRT1 line were responsive, indicating cell-type specific roles such as repressing glucose-induced insulin secretion in β-cells; thus, showing no major effects in HEK293T cells^34^ **(Supplementary Fig. 2A)**. Further, we observed that only 2.1% (737/35,245) of the responsive fragments in the FOXA1 line and 22% (4,575/20,987) in the ATF2 line overlapped with their respective binding sites identified through ChIP-seq. These results suggest that most responsive fragments are likely bound by other TFs that compensate for the loss of ATF2 or FOXA1, or alternatively, these could reflect stress response mechanisms. In fact, motif enrichment analysis of these responsive fragments revealed enrichment for other TF motifs, such as GATA factors, HOX factors, KLF factors and P53 family of TFs.

We also observed that the activity of individual responsive fragments varied across the KO lines suggesting that enhancer “state” reflects both sequence and context **(Figure 3A and 3B**). For instance, an active fragment (chr17:80693000-80695800) in CC, showed increased activity in FOXA1, TCF7L2, and SCRT1 lines, but had decreased activity in CTCF, ATF2, and LEF1 lines. Therefore, the same sequence can manifest distinct quantitative enhancer functions across TF KO lines. In fact, modality-specific assays such as mSTARR-seq have shown that responsiveness can be conditional on additional molecular context such as DNA methylation state of enhancers^35^. To further dissect these trends, we calculated the variance in activity for each fragment across all lines and observed a strong correlation with their native activity in the CC line, i.e., highly active fragments in CC showed highly variable activity across the KO lines (Pearson r=0.56) **(Fig. 3C)**. For example, the fragments located on chr2:19228797–19229297 and chr16:89995593–89996093 showed the highest variance in activity across lines and were both annotated as active enhancers in CC. However, the chr2 fragment was repressed in ATF2 and LEF1 lines but induced in FOXA1 and SCRT1 lines, whereas the chr16 fragment was strongly induced in ATF2 and FOXA1 but lost activity in the CTCF line. These observations suggest that the regulatory potential of candidate enhancer fragments is dependent on the cellular regulatory state, in particular TF availability. To investigate this further, we divided the fragments into deciles based on their activity in CC and calculated the percentage of responsive fragments for each KO line within these deciles **(Fig. 3D)**. We found that the percentage of responsive fragments and the number of TF binding motifs increased exponentially across these deciles **(Fig. 3D and 3E)**. These results suggest that motif “hotspots” not only drive higher baseline activity but also modulate across different TF KO lines, reflecting variation in cellular contexts. This observation is consistent with large-scale studies using synthetic and genomic libraries showing that motif content and combinatorial motif load correlate with enhancer activity^36–39^. However, the percentage of responsive fragments in the lowest CC activity decile was significantly higher in the ATF2 KO line compared to the others **(Supplementary Fig. 2B).** For instance, chromosomal fragment chr8:6,649,383–6,649,883 located in the intronic region of *MCPH1* showed a 23-fold increase with ATF2 KO while being inactive in CC **(Fig. 3A)**.

**Figure 3:**
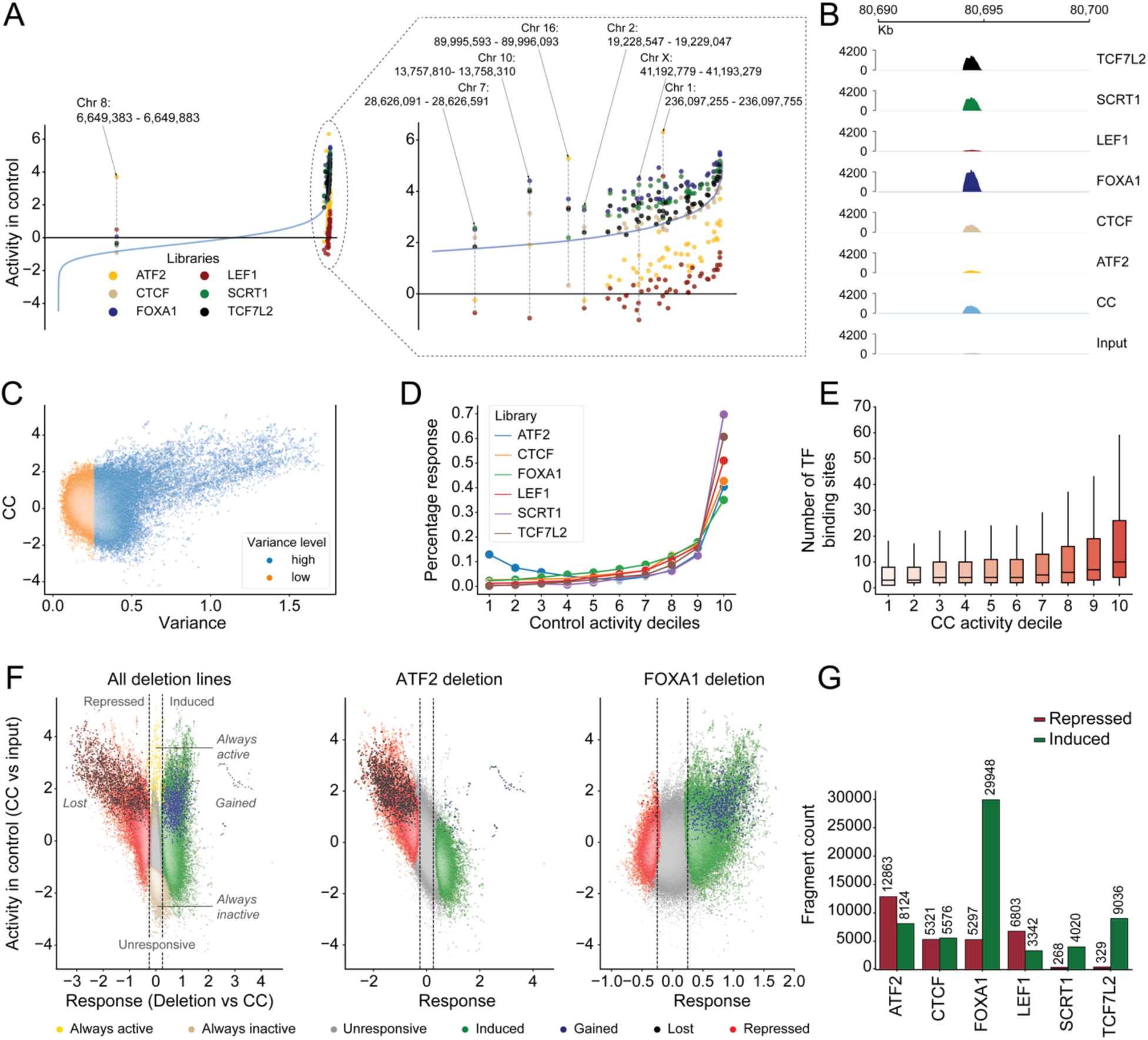
Assaying enhancer fragment activity and comparing response. **(A)** Activity of selected fragments with high variability across TF KO lines compared to activity in CC. The blue line denotes the activity observed in CC. Fragments which have variable activity in the KO lines and are highly active in CC are highlighted in the box. **(B)** Variation in the number of normalized reads observed in a representative fragment on chr17:80690000-80700000 across all input, CC, and all KO lines. **(C)** Variance in response (activity in KO/CC) for each fragment compared to activity in CC. **(D)** Percentage of fragments showing response (adjusted *P*<0.01) in each TF KO line across the CC activity deciles. **(E)** Box plot with the number of TF binding sites observed across each CC activity decile. **(F)** Activity map showing response of fragments to TF KO versus activity in CC highlights different categories of enhancers. Definition of fragments, (a) *Unresponsive*: Fragments with insignificant change of activity in KO lines compared to CC, (b) *Always active*: Fragments unresponsive in CC but showed enhancer peaks in all KO lines, (c) *Always inactive*: Fragments unresponsive in all lines, (d) *Induced*: Fragments with significant increase in activity in the KO line compared to CC, (e) *Repressed*: Fragments with significant decrease in activity in the KO line compared to CC, (f) *Gained*: Fragments with no peaks in CC but showing enhancer peaks in the KO lines, (g) *Lost*: Fragments showing enhancer peaks in CC but no peaks at all in the KO lines. **(G)** The number of induced and repressed fragments observed for each KO line is shown.

Next, we classified responsive fragments as KO “induced” or “repressed” **(Fig. 3F and 3G),** based on whether the enhancer activity was significantly increased or decreased in the KO lines relative to CC. All KO lines showed both induced and repressed fragments, although in varying proportions, suggesting both activator and repressor roles for these TFs. While most responsive fragments were induced in the FOXA1 (85%), SCRT1 (94%) and TCF7L2 (96%) lines, a majority of responsive fragments in ATF2 (61%) and LEF1 (67%) lines were repressed **(Fig. 3G)**. In fact, 67% of the repressed fragments in LEF1 line were also repressed in ATF2 line, suggesting that these two TFs may share activating roles in convergent biological pathways (**Supplementary Fig. 2C**). Further, we found a subset (0.7%, 1,192/251,256) of fragments that were inactive in CC but significantly induced (i.e., gained activity) with TF deletion, and conversely fragments (81%, 1,635/2,376) that were active in CC but significantly repressed (i.e., lost activity) with TF deletion **(Fig. 3F, Supplementary Fig. 2D)**.

### Effect of differential enhancer activity on target gene expression

Next, we assessed how differential enhancer activity influenced target gene expression by focusing on induced enhancers driving upregulation and repressed enhancers leading to downregulation of genes. Using a combination of the ‘nearest-gene method’^7^ and a modified ‘activity-by-contact’ model^13^ (see **Methods**), we found that 84% (47,422 of the 56,617) of differentially active enhancers identified across all libraries mapped to their target genes (**Supplementary Data 3**). For instance, depletion of ATF2, a known transcriptional activator (**Table 1**), led to widespread repression of enhancers enriched for motifs from both the ATF and TP53 families. Further, these repressed enhancers resulting from ATF2 KO targeted signaling pathways such as Wnt and Hippo, leading to downregulation of genes such as *MAPK10*, *TCF7L2*, *CREBBP*, *LEF1*, and *BMP7* **(Fig. 4A)**. Previous studies have linked Wnt signaling genes such as *LEF1* and *TCF7L2* to ATF2 family of TFs^40^. Consistent with this finding, LEF1 KO predominantly led to repression of enhancers, with substantial overlap of lost enhancers between the ATF2 and LEF1 lines (**Supplementary Fig. 2C**). In fact, we identified sixteen such enhancer fragments in the ATF2 line whose loss of activity led to downregulation of *THRB*, *TCF7L2*, *KMT2E*, *STK17A*, and other genes involved in p53 signaling **(Supplementary Fig. 3A)**. Enrichment of the p53 motif among repressed enhancers was exclusively observed in ATF2 and LEF1 KO libraries, supporting the convergence of these transcription factors on a common regulatory network. Notably, our framework also recovered known regulatory connections. For example, ATF2 KO revealed a gained enhancer linked to upregulation of *PSEN1*, a known transcriptional target of ATF2 in the Notch pathway^41^. Similarly, CTCF KO resulted in the loss of enhancer activity at the *FIRRE* locus, which harbors multiple CTCF binding sites, leading to reduced expression of the *FIRRE* lncRNA^42^ **(Fig. 4A)**.

**Figure 4:**
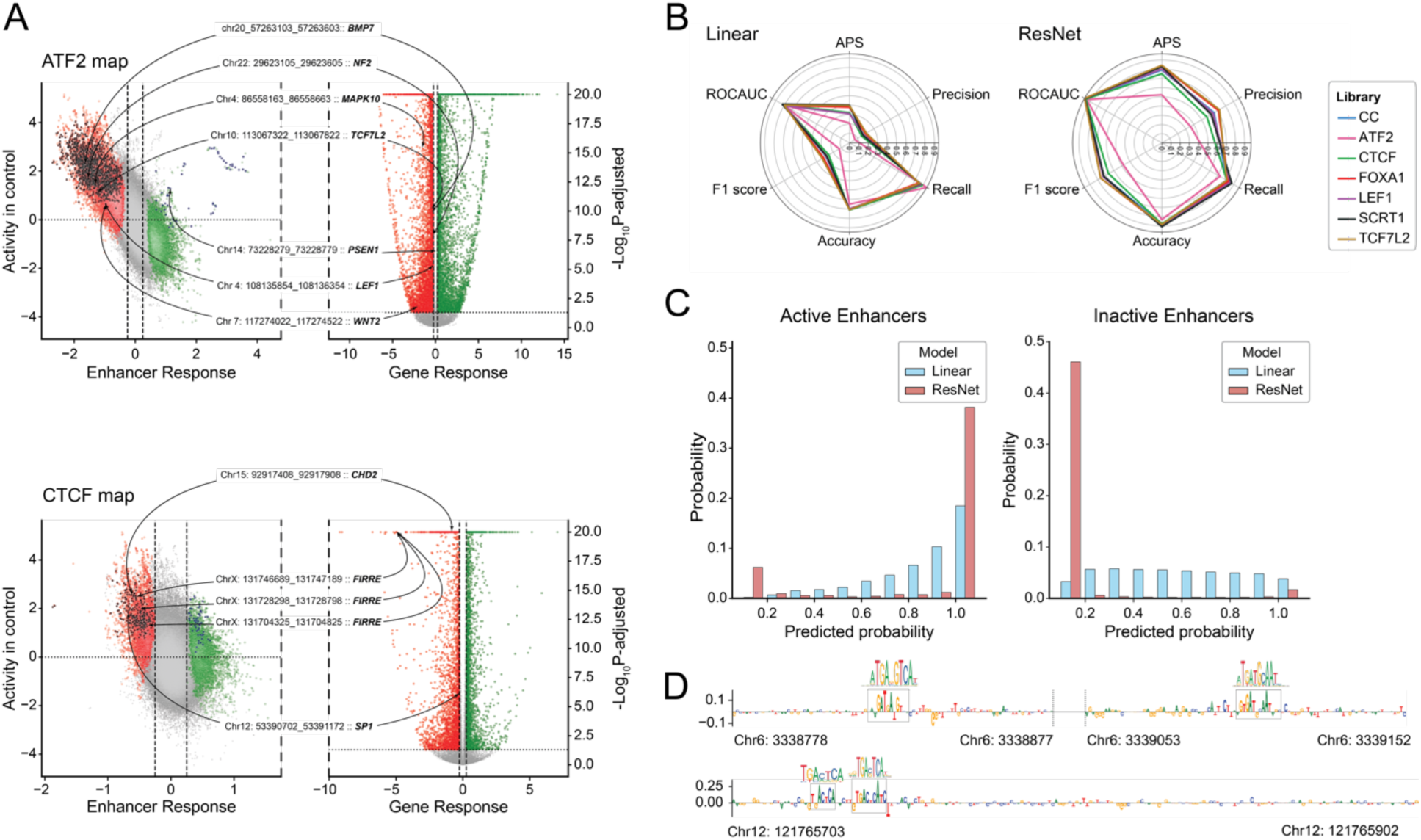
Mapping enhancers to target genes and learning the regulatory code. **(A)** Activity maps for differential enhancer activity and volcano plot for differential gene expression showing specific enhancers mapped to their target genes in the ATF2 KO cell line (top) and the CTCF KO line (bottom). **(B)** Radar plots showing the enhancer activity prediction metrics for the linear model (left) and ResNet model (right). **(C)** Probability of predicting active enhancers in high STARR-seq coverage regions (left) and inactive enhancers in low STARR-seq coverage regions (right) using linear and ResNet models. The probability distributions showed a rightward shift for high coverage regions and a leftward shift for non-peak fragments under the deep learning model. **(D)** Motif combinations highlighted by ResNet model while predicting enhancer activity.

In contrast, depletion of FOXA1 induced enhancer response leading to predominantly gained enhancers (**Fig. 3F**), and upregulation of genes involved in protein dephosphorylation (such as *PTPA*, *ENSA*, and *CAMTA1*) and RNA polymerase II–mediated transcriptional regulation (such as *SMARCB1*, *OTUD7B,* and *CHD2*) **(Supplementary Fig. 3B)**. FOXA1 functions as a pioneer factor that primes enhancers and promotes RNA polymerase II engagement through the Mediator–Cohesin axis^43–45^ (**Supplementary Table 1**). The upregulation of RNA polymerase II regulatory genes by gained enhancers following FOXA1 KO suggests a compensatory activation of a polymerase-coupled remodeling program through chromatin remodelers such as *CHD2* and *SMARCB1*. Overall, our framework identifies TF-specific responsive enhancers that regulate expression of their target genes.

### Using deep learning models to predict enhancer activity

A combined analysis of UDI-UMI-STARR-seq and RNA-seq data suggested that specific TF binding sequence motifs on the enhancer fragments defined their patterns of activity for each KO line. This observation is in line with previous MPRA studies using base-level resolution deep learning models that identified specific sequences predicting enhancer activity^14,36,46^. Termed as enhancer grammar, these predictive rules encoded by the sequences can range from additive effects where each relevant TF motif independently contributes to activity with relatively weak constraints on their arrangement to context-dependent motif syntax requiring specific motif combinations, spacing, or orientation for activating enhancers. To this extent, we trained a linear classifier and a deep learning model based on a residual network (ResNet) architecture^47^ to predict enhancer activity across all KO lines (**Methods**). The ResNet models consistently outperformed the linear baselines across all evaluation metrics, indicating that context-dependent motif syntax plays a crucial role in governing enhancer activity in the KO lines **(Fig. 4B)**.

We evaluated the relationship between model predictions and experimentally measured enhancer activity by examining the distribution of predicted enhancer activity probabilities across all test fragments. We found that the deep learning model consistently assigned higher probabilities (0.9–1.0) to a greater proportion of identified enhancer peaks than the linear model, while conversely assigning lower probabilities (0.0–0.1) to a greater proportion of non-peak fragments **(Fig. 4C)**. Further, STARR-seq fragments with higher sequence coverage showed higher probabilities (>0.9) from the ResNet model than from the linear model (Fisher’s exact test, OR=1.85, 95% CI=1.69, 2.03, *P*=3.63 × 10^−41^), indicating that the deep learning framework is better calibrated to distinguish experimentally validated enhancer activity from background signal, possibly due to its ability to learn non-linear patterns of motif syntax that the linear model cannot capture. To investigate the sequence determinants driving this improved performance, we applied Integrated Gradients **(Methods)**, a feature attribution method that identifies informative nucleotide positions contributing to model predictions. Aggregating these attributions using TF-MoDISco^48^ revealed sequence motifs that were common between the linear and ResNet models for predicting high-confidence peaks across libraries **(Supplementary Fig. 4A)**. We also observed recurrent motif combinations uniquely captured by the ResNet model in high-coverage regions that were not confidently predicted by the linear model across multiple KO lines. *For example*, we observed adjacent motifs for JUN and FOSl1 on chr12 and for ATF2 and CEBPG on chr6 within active enhancer regions of the FOXA1 KO library **(Fig. 4D)**. These TF pairs are known to dimerize and modulate DNA-binding specificity and transcriptional activity^49–51^. Similarly, a CREB1 motif was detected in proximity to an EPAS1 motif on chr1, suggesting potential regulation of EPAS1 transcriptional targets by CREB1 **(Supplementary Fig. 4B)**. Other instances where combinations of adjacent motifs led to high-confidence enhancer predictions include adjacent ATF2 and ATF4 motifs at chr2:45953616 and chr5:136042862 in the CC line **(Supplementary Fig. 4B)**. Notably, the differences in spacing between these motifs suggest that linear distance does not constrain enhancer activity, consistent with the concept of soft motif syntax^14^. Additionally, in several cases, the attribution scores extended into flanking sequences adjacent to the core motifs, consistent with prior observations of context-dependent enhancer recognition by deep learning models^36^ **(Supplementary Fig. 4C)**.

### Regulatory network for neurodevelopmental disease-associated 16p12.1 deletion

We next sought to assess the effect of 16p12.1 deletion on enhancer activity and consequent transcriptome changes. The approximately 500 kbp 16p12.1 deletion has been associated with a range of neurodevelopmental outcomes, including intellectual disability/developmental delay, autism, depression, anxiety, and schizophrenia^52,53^. Although analyses of *Drosophila* and human induced pluripotent stem cell (iPSC)-derived neuronal models have implicated multiple pathways affected by the deletion, the regulatory networks connecting these pathways remain unclear.

To address this, we generated the deletion using the CRISPR/Cas9 strategy and performed STARR-seq in parallel with RNA-seq (**Fig. 5A**, **Supplementary Information).** Comparing enhancer activity in the 16p12.1 deletion line with CC revealed that 2% of the fragments were responsive, with 59% (3,147/5,330) of these fragments belonging to the higher activity deciles in CC **(Fig. 5C)**. Linking enhancer activity to expression of putative target genes revealed that gained enhancers were associated with upregulation of genes involved in axon guidance and synaptic plasticity (such as *PRKCA*, *GNAI1*, and *ROBO1*) as well as brain morphology and connectivity (such as *RUNX2* and *ROBO1*) (**Fig. 5D**). Independent studies have identified mutations in these genes to be associated with congenital anomalies and developmental disorders^54–57^, highlighting their potential contributions to the phenotypic spectrum observed in individuals with the 16p12.1 deletion. In contrast, lost enhancers were associated with downregulation of *CTIF*, a CBP80/20-dependent translation initiation factor involved in the pioneer round of translation and nonsense mediated decay^58^. More broadly, perturbation of translational control and mRNA surveillance has been implicated in neurodegenerative risk, providing a plausible mechanistic bridge between enhancer loss and neuronal vulnerability^59,60^. Pathway analysis revealed that genes upregulated by gained enhancers were enriched for PI3K-AKT (including *CDKN1A*, *LPAR6*, PRLR) and MAPK (*KITLG*, *PRKCA*) signaling, mirroring findings from our studies using neural progenitor cells^61^. Deep-learning models trained on the 16p12.1 deletion library recapitulated trends observed in other KO datasets, with ResNet outperforming linear models across performance metrics (**Fig. 5E**). Model interpretation and motif enrichment analysis converged on p53 and ATF/bZIP family motifs as major determinants of enhancer activity, suggesting that stress-responsive transcription factors occupy a substantial fraction of active elements in this context (**Fig. 5F**). Since our study was performed in a cancer cell line, we note that expression patterns of nervous system genes may differ considerably from those in neuronal cells. However, our findings demonstrate the utility of this approach for uncovering downstream effects of genomic variants on regulatory networks and cellular function.

**Figure 5:**
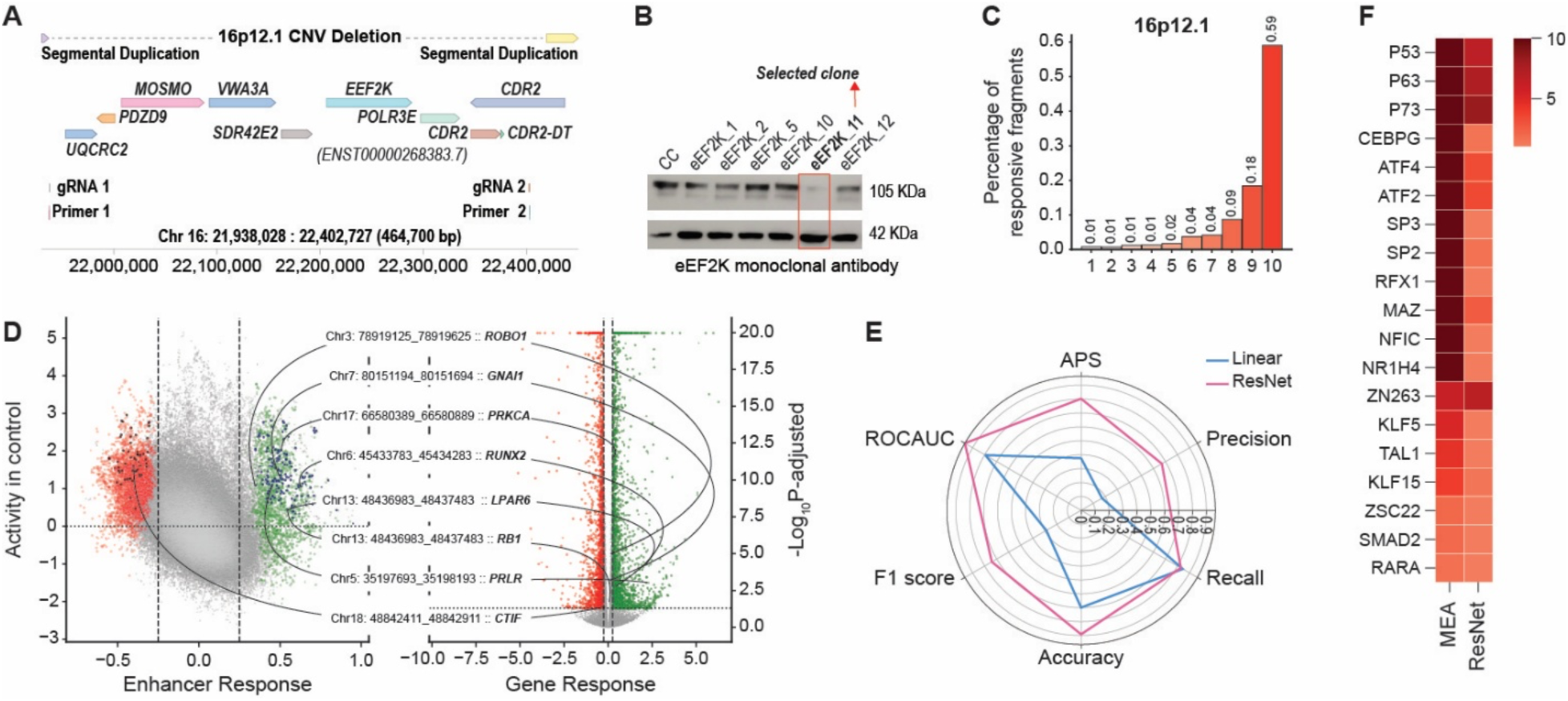
Applying UDI-UMI-STARR-seq and RNA-seq to assay the 16p12.1 deletion. **(A)** The chromosomal location of the 16p12.1 region and its constituent genes. **(B)** Western blot for the eEF2K protein to validate the deletion. **(C)** Percentage of responsive enhancers due to the 16p12.1 deletion across the CC activity deciles. **(D)** Enhancer activity and differential gene expression map of the 16p12.1 deletion with induced and repressed fragments targeting differentially expressed genes. **(E)** Evaluation metrics of the linear and deep learning models predicting enhancer activity. **(F)** Common motifs enriched within the enhancer peaks of the 16p12.1 deletion library as well as those identified by the ResNet model to predict enhancer activity.

## DISCUSSION

In this study, we developed a framework that systematically measured the downstream effects of genetic mutations on enhancer activity and gene expression. Several themes have emerged from our study. *First*, genetic perturbations, such as TF or a pathogenic CNV deletion, can reshape enhancer activity at a global scale revealing coordinated patterns of induced and repressed enhancers that regulate downstream transcription. While previous STARR-seq studies have largely focused on allele-specific effects of single-nucleotide variants with modest, context-limited outcomes^18,62–66^, our results show that high-effect variants have a relatively larger outcome on enhancer activity. Comparable global shifts in enhancer activity have been observed following hormone or environmental stimulation, where glucocorticoid or retinoid signaling triggers genome-wide enhancer activation and repression^8,17^. *Second*, enhancer responses to a specific genetic perturbation are not uniform but instead fall along a spectrum, and is skewed towards either induced or repressed enhancers depending on the perturbation. *For example*, FOXA1 and TCF7L2 depletion led to primarily induced enhancers, whereas ATF2 and LEF1 depletion resulted in widespread repression. Similar asymmetries were reported in a previous TF-centric enhancer perturbation study, where cofactor balance and motif grammar determined whether regulatory element activity is induced or repressed due to a perturbation^67^. The observation that the same enhancer sequence can switch between activation and repression depending on the perturbation highlights the context-dependent and combinatorial nature of enhancer regulation. Our results mirror recent findings on dual function regulatory elements where the same region can act as either an enhancer or a silencer depending on the cellular context^68–70^. *Third*, enhancer fragments showing the strongest response overlapped with multiple motif hotspots, that potentially act as super enhancers, consistent with their increased sensitivity to perturbation. Previous studies have shown that TFs colocalize at these hotspots to remodel chromatin and recruit coregulators necessary for transcriptional activation^71,72^. The concentration of perturbation-responsive enhancers within such domains underscores their central role as hubs for dynamic control of gene expression and cellular state transitions. *Fourth*, the deep learning-based sequence-to-function models indicate that enhancer activity is governed not merely by motif presence but by higher-order motif syntax and flanking sequence context. Deep learning frameworks such as BPNet and DeepSTARR have shown that motif multiplicity, spacing, and orientation encode quantitative enhancer activity^14,36^. Consistent with these observations, our models identified recurrent motif pairs with flexible spacing and orientation preferences, and demonstrated that inclusion of adjacent nucleotides flanking TF motifs improved predictive accuracy. These findings reinforce the notion that enhancer logic is best described by probabilistic grammar rather than discrete motif presence. *Finally*, we observed that linking differentially active enhancers to differentially expressed target genes increases the resolution of measuring perturbation effects by adding a second modality beyond transcriptomics. Our multi-modal analysis revealed convergent mechanisms between ATF2 and LEF1 on p53-linked transcriptional programs and compensatory, chromatin remodeling following FOXA1 depletion. Consistent with this, paired chromatin accessibility and transcription has repeatedly outperformed transcriptomics-only study designs in recovering regulatory drivers under perturbation or state change. For example, multimodal enhancer-gene maps integrating bulk or single-cell ATAC- and RNA-seq datasets have improved TF-to-target-gene inference^73–75^, and recent perturbation or activation studies have identified key regulatory elements only when accessibility and expression are measured together^76,77^. These results demonstrate the utility of our framework to capture downstream effects of a perturbation through enhancer mediated regulatory mechanism. Further, by applying this framework to an NDD associated CNV deletion, we demonstrate its generalizability to any genetic perturbation.

Although effective, our approach is limited by the choice of cell line. While a cancerous cell may not be ideal to test the effects of a mutation that might have neuronal or other disease related consequences, as a proof-of-concept study we demonstrate that our framework can be applied to any other context depending on the subject of the research. Moreover, our assay measures enhancer activity in an episomal context and may not completely capture endogenous effects such as local chromatin state, genomic position, or three-dimensional contacts. We selected this approach to obtain an unbiased readout of enhancer activity unlike in endogenous assay designs such as LentiMPRA where the target fragments are embedded into the genome at random and the cumulative activity of each sequence is measured using unique barcodes^78^. By avoiding random genomic integration, our assay minimizes confounders such as locus-dependent positional advantages and competition from neighboring regulatory elements. Additionally, connecting differentially active enhancers to their target genes is an area of active research. The target genes identified by our method through predictive models such as the ABC model may not be completely accurate. Therefore, experimental validations in the specific context are required. Finally, using a targeted library allowed us to measure enhancer activity at nucleotide resolution. However, it also restricted our analysis to regions with strong prior evidence of enhancer potential rather than providing genome-wide coverage, an approach that can be complemented by future genome-wide libraries.

## METHODS

### STARR-seq plasmid library preparation

The STARR-seq plasmid library was prepared as described previously^24^ with modifications derived from CapSTARR-seq^21,63^ and UMI-STARR-seq^22^ protocols. A brief description of major steps is provided below.

#### (A) Shearing and fragment size selection

Human genomic DNA (catalog #G3041, Promega, WI, USA) isolated from multiple anonymous male and female donors was sheared to an average fragment size of 500 bp using a Covaris sonicator (Covaris, MA). Fragment size selection was performed using Blue Pippin (Sage Sciences, MA, USA) in five replicates, and the size distribution was validated using TapeStation (Agilent Technologies, CA, USA). DNA shearing, size selection and validation was performed at the Genomics Core Facility, Huck Institutes of Life Sciences, Pennsylvania State University.

#### (B) Insert preparation

Equal amounts of size-selected DNA fragments (by weight) were pooled and used for subsequent library preparation in three replicates following the Kapa HyperCap workflow (Roche, IN, USA). Briefly, fragments were end-repaired and dA-tailed in triplicates followed by ligation of custom designed oligonucleotides containing Illumina read 1 and read 2 adapters. Cloning overhang sequences were added using ligation mediated PCR (LM_PCR) for nine cycles. PCR amplification was performed using Kapa HiFi HotStart ReadyMix (catalog #KK2602, Roche, IN, USA), and adapter ligation efficiency was validated by assessing the expected fragment length increase at each step using a TapeStation.

To enrich the library for our target regions, we performed hybridization and capture using custom designed SeqCap EZ XL probes (catalog #08247510001, KAPA Biosystems, MA, USA and Roche, IN, USA). Three independent captures were carried out according to manufacturer’s protocol each starting with 1µg of purified LM_PCR product per capture. The captured DNA was pooled in equal proportions prior to subsequent library amplification. COT Human DNA (catalog #11581074001, Sigma Aldrich, MO, USA) was used to prevent non-specific binding of probes to repetitive genomic regions, and custom designed blocking oligonucleotides were used to prevent sequencing adapters from binding to each other during hybridization. All purification steps were performed with Ampure XP beads (catalog #A63881, Beckman Coulter, USA), using bead-to-sample volume ratio of 1.8X (adjusted for 500 bp fragments). The final amplified insert pool was purified twice and eluted in 40 µl, yielding concentrations >25 ng/µl and 260/280 and 260/230 ratios of 1.8 and 2.2, respectively, indicating high quality DNA. Quality control metrics for all intermediate steps are provided in **Supplementary Information**.

#### (C) STARR-seq plasmid library cloning

The human STARR-seq vector (addgene #99280) containing the ORI sequence^79^ was digested with restriction enzymes SalI-HF (catalog # R3138S, New England Biolabs, MA, USA) and AgeI-HF (catalog #R3552S, New England Biolabs, MA, USA), as previously described^22,24^. We performed 24 independent digestion reactions with 1 µg of vector per reaction. Reactions were resolved on a 1% agarose gel using electrophoresis. The digested backbone at approximately 2-kbp mark was extracted using Zymoclean Gel DNA Recovery Kit (catalog #D4001, Zymo Research, CA, USA). Up to eight purified samples were pooled and re-purified using a DNA clean and concentrator kit (catalog #D4003, Zymo Research, CA, USA) followed by 1X Ampure XP bead purification to maximize product purity. Final products yielded DNA concentrations >35 ng/µl in 30 µl with purity ratios (260/280 and 260/230) between 1.8-2.2.

Assembly reactions were performed using NEBuilder HiFi DNA assembly mix (catalog #E2621L, New England Biolabs, MA, USA) with a 2:1 insert-to-vector molar ratio. The total molarity of each reaction (vector + insert) was maintained between 30 to 200 fmol in accordance with the manufacturer’s protocol. 24 independent assembly reactions were carried out and up to four reactions were pooled and further purified using 1X Ampure XP beads or DNA clean and concentrator kit and eluted in 15µl of water. The pooled and purified reactions were transformed into NEB5-alpha (catalog #C2989, currently discontinued) and NEB10-Beta (catalog #C3020K, New England Biolabs, MA, USA) electrocompetent cells. For each transformation, 3 µl of purified and cloned library was mixed with 25µl of electrocompetent cells and electroporated using two pulses at 1800V. Subsequently, the mix was transferred to 972 ml of pre-warmed SOC media and incubated at 300 rpm at 37⁰C for 1 hour.

Transformation efficiency was estimated by calculating colony forming unit (CFU) per µg of DNA using a supercoiled spCas9 plasmid (catalog #62988, addgene, MA, USA) as positive control. To understand cloning efficiency, we performed serial dilution of the transformed mix. 100 µl of the SOC mix from each transformation reaction was serially diluted in a 1:10 ratio, six times, in 900 µl LB media per dilution. 100 µl of each dilutant was spread on an ampicillin containing LB plate (100 µg/ml) and incubated overnight. The remaining 900 µl of transformation reactions were added to 150 ml to 250 ml LB broth cultures (to a total 3 liters) and incubated overnight for 16 hours at 37⁰C and 300 rpm.

#### (D) Library isolation, storage and validation

Following incubation, we ensured that the cultures reached an optical density of approximately 2.6 prior to library isolation. From each culture, 5 ml of culture broth was reserved to generate glycerol stocks of the library for storage. Plasmid DNA was isolated from the remaining culture volumes using Zymopure II plasmid Maxiprep kit (catalog #D4203 Zymo Research, CA, USA), according to manufacturer’s instructions. Individually isolated samples were pooled by equal mass, treated for endotoxins using columns provided with the kit, and normalized to a final library concentration of 38.5ng/µl. To assay library coverage and complexity, we first PCR amplified the library in triplicates using Q5 Hot Start High-fidelity 2X master mix (catalog #M0494L, New England Biolabs, MA, USA) and NEB Multiplex Oligos, primer set 1 (catalog # E7335S, New England Biolabs, MA, USA), with indexes 4, 6, and 12 based on the manufacturer’s protocol. Next, libraries were pooled and sequenced using Miseq nano 150 bp paired end sequencing at Huck Genomics Core Facility.

### Cell culture

HEK293T cell (catalog #CRL-3216, American Type Culture Collection, VA, USA) culture was carried out according to ATCC guidelines in Dulbecco’s Eagle Modified Media (DMEM) premixed with 4500 mg/L glucose, L-glutamine, sodium pyruvate and sodium bicarbonate (catalog #D6429, Sigma Aldrich, MO, USA). DMEM was additionally supplemented with 10% Fetal Bovine Serum (FBS) (catalog #F2442, Sigma Aldrich, MO, USA), 1% Penn/Strep (catalog #4333, Sigma Aldrich, MO, USA), 1% HEPES (catalog #H0887, Sigma Aldrich, MO, USA) and 1% MEM Non-essential amino acids (catalog #M7145, Sigma Aldrich, MO, USA). Cells were cultured at 37⁰C and 5% CO_2_ within a sterile incubator . All transfections were performed using Lipofectamine 3000 (catalog ##L3000008, Thermo Fisher scientific, MA, USA) according to scaled protocols following the manufacturer’s instructions. Stable CRISPR/Cas9 mediated deletions were generated using a dual gRNA system and validated using western blot (**Supplementary Information**).

### STARR-seq screening

STARR-seq screening libraries were generated as described previously^24^. A brief overview of major steps is provided below.

#### (A) Library transfection and RNA Isolation

The STARR-seq library was independently transfected into wildtype HEK293T cells and each knock-out line in three replicates at approximately 1 µg library per 1 million cells (approximately 38 million cells on 15 cm dishes). Total RNA was isolated 24 hours post transfection using the TRIzol Plus RNA purification kit (catalog #12183555, Thermo Fisher scientific, MA, USA) with 6 ml of TRIzol per 15 cm dish. To assess sample quality, RNA Integrity Analysis (RIN) was measured on a TapeStation (Agilent Technologies, Santa Clara, CA), with RIN scores >9.5/10 for all samples (data available upon request). mRNA was isolated using Dynabeads mRNA Purification kit (catalog #61006, Thermo Fisher scientific, MA, USA), according to the manufacturer’s protocol. In brief, each replicate was diluted to ∼750 ng/µl and used up to 75 µg of total RNA per purification reaction with beads reused up to five times for the same sample replicate. For each replicate, 10-12 isolation reactions were performed, and all reactions were pooled, recovering up to 25 µg of mRNA per replicate. Residual DNase was removed by treating mRNA with TURBO DNase (catalog #AM1907, Thermo Fisher scientific, MA, USA) and samples were further purified using RNAClean XP beads (catalog #A63987, Beckman Coulter, USA).

#### (B) cDNA library preparation

For each replicate, 12.5 µg mRNA (2.5 µg per reaction) was reverse transcribed to first strand cDNA using Superscript III reverse transcriptase (catalog #18080044, Thermo Fisher scientific, MA, USA) and a STARR-seq reporter transcript specific primer^22,24^. Replicate reactions were pooled, residual RNase was removed using RNase A and samples were further purified using 1.4X AMPure XP beads. Second strand synthesis was performed using another STARR-seq reporter transcript specific second strand synthesis primer (**Supplementary Table 2**) and Kapa HiFi HotStart ReadyMix in a single cycle PCR reaction followed by product purification using 1.4X AMPure XP beads.

#### (C) UMI addition and junction PCR

Unique Molecular Identifiers (UMI) and sequencing indexes were added using custom i7-UMI-P7 primers containing 10 bp UMIs in between 8 bp i7 index barcode and P7 adapter sequence **(Fig. 2B)**. Unlike the UMI-STARR-seq protocol suggested by Neumayr et al.^22^, the i7 index barcode was retained to enable unique dual indexing (along with i5 index) and reduce index hopping in our libraries during sequencing^23^. The final number of sequencing cycles was adjusted accordingly. Following each step, the sample was purified with 1.4X AMPure XP. Junction PCR (jPCR) was then performed (5 reactions per replicate) using jPCR primers. Reactions from each replicate were pooled and purified using 0.8X AMPure XP beads.

#### (D) STARR-seq sequencing library preparation

##### (a) Output sequencing library

For each output library, 10–20 µl of jPCR product was used as a template for the final amplication step, a low-cycle sequencing ready PCR. Reactions were performed for five cycles with a standard Illumina i5 primer, containing an i5 index barcode, and a custom P7 primer, containing only the P7 adapter sequence, using Kapa HiFi HotStart ReadyMix. All products were assessed by agarose gel electrophoresis to confirm fragment-size distribution and check for overamplification. Samples were then purified with 1X AMPure XP beads.

##### (b) Input sequencing library

To generate the input library, we performed a single-cycle PCR amplification of the plasmid library in three replicates under conditions analogous to the UMI-addition PCR for the output libraries, except that we included a custom i7-UMI-P7 primer together with the i5 primer to introduce both indexes simultaneously^24^. This ensured that the *input* and *output* sequencing libraries both shared the same sequence architecture. PCR products were purified with 1X AMPure beads and then further amplified using the i5 and P7 primers for five cycles, similar to the output libraries. Subsequently, fragment-size was verified by agarose gel electrophoresis, residual vector backbones were removed by excising the band of correct length, and the DNA was isolated using Zymoclean Gel DNA recovery kit. The recovered DNA was further purified using 1X AMPureXP beads and eluted in 25 µl of DEPC water. Overall, we generated 24 output and 3 input sequencing libraries.

### Library sequencing

Libraries were sequenced on an Illumina NextSeq 2000 sequencer. All 27 libraries were quality-assessed on a TapeStation and pooled in an equimolar fashion. The library pool was then diluted and mixed with PhiX and loaded onto the sequencer. Each library was sequenced to approximately 45 million reads (1.2 billion reads in total) using 150 bp paired end reads. Sequencing was carried out for 18 cycles for i7 and 8 cycles for i5. The first 8 nucleotides from i7, i.e., the index barcode sequence, were used for parsing and demultiplexing the libraries from the pooled run.

### Candidate enhancer library selection

We curated candidate enhancer regions from ENCODE ChIP–seq datasets generated in HEK293 or HEK293T cells. ChIP-seq sites for the enhancer-associated histone marks H3K27ac and H3K4me1 were merged with BEDTools^80^. To specifically focus on distal enhancer elements, sites within 2 kbp upstream and downstream of annotated Transcription Start Sites (TSS) were removed. The distal set was then intersected with ChIP-seq sites for 226 transcription factors (TFs) to obtain regions marked by both enhancer-associated histone modifications and TF binding, while excluding regions bound by a single TF. In addition, due to limited HEK293/HEK293T data for certain factors, we supplemented the set with high-confidence enhancer regions defined by p300, CHD8, and SMARCA4 ChIP–seq in K562 and HeLa cells. The final set comprised 46,142 regions spanning 32.9 Mbp of the human genome, with individual ChIP–seq intervals ranging from 170 bp to approximately 10 kbp.

### STARR-seq reads processing and data quality assessment

#### (A) Read deduplication, alignment, and filtering

Raw FASTQ files of STARR-seq libraries obtained directly from the sequencer were demultiplexed using STARRDUST (STARR-Seq Deduplication on UMI Sequence Tags), an in-house tool tailored to the unique library design where UMIs were placed at both the 5’ and 3’ ends of each amplicon. Typically, programs such as UMI-tools or UMI-dedup are used to deduplicate mapped reads from Sequence Alignment Map (SAM) files. However, depending on the UMI strategy, and sequencing assay, deduplication prior to read mapping is also a feasible strategy when combined with quality filtering steps. STARRDUST performs pre-alignment quality control and UMI deduplication in a single pass. Specifically, it filters (1) all reads with Q-Scores less than 30, (2) reads containing unresolved “any” base (N) especially in the designated UMI regions of the reads, and (3) duplicated inserts containing the same 5’ and 3’ UMI-pair. Reads with swapped i5 and i7 indexes are filtered out during the demultiplexing step due to known apriori pairing of i5 and i7 indexes that is unique to each of the samples that were sequenced. For deduplication, the probability of the same amplicon getting the same 5’ and 3’ UMI-pair twice, from the PCR step where UMIs are added, is assumed to be a very low, and thus provides a simple, yet consistent basis for eliminating both PCR and optical duplication without the need for aligning the reads. To prevent deduplication from becoming overly stringent, however, we do not filter out near-duplicate inserts with same UMI-pairs, especially because deduplication is performed after the quality-filtering step. Thus, we only expect a few real duplicated reads to survive our filtration strategy, those that occur due to systemic base-calling errors. This also makes the deduplication strategy computationally efficient, since we only need to maintain a dictionary of the actual UMI-pairs with only the message digest of the sequenced reads, for rejecting reads, as we iterate through a pair of FASTQ files.

Following de-multiplexing, the reads were aligned to the human reference genome version GRCh38 using BWA-MEM with default parameters^81^. Low quality, multi-mapped and off-target reads were filtered using Samtools^82^ with the following parameters: -F 2828 -f 2 -q 30. The remaining reads were used in downstream analysis.

#### (B) Data quality assessment

To assess library data quality, we calculated reads per kilobase million (RPKM) normalized read counts across all regions of interest for each library replicate and examined the distribution of read counts per region within and across libraries. Reproducibility was measured by calculating Spearman correlation of output-over-input normalized read fold changes between library replicates.

### STARR-seq peak calling and peak quality assessment

To identify active enhancers, input and output library replicate bam files were merged and library peaks were identified using STARRPeaker^26^ with default parameters. To measure the reproducibility of peaks between library replicates, spearman correlation of output-over-input normalized read fold changes of the peaks was calculated. To compare activity between peaks and exonic regions, library regions which overlapped with annotated exons from the reference human genome (GRCh38) were first identified. Subsequently, output over input fold change of normalized reads was calculated for both peaks and the identified exonic regions across all libraries. Finally, the fold change distributions between peaks and exonic regions were compared using t-test.

To assess concordance between peaks and open chromatin regions, DNase-Seq dataset of HEK293T cells were obtained from ENCODE database. The filtered bam files from Input and CC library were converted to bigwig format using deepTools^83^. Finally, the normalized read counts from DNase-Seq and output-over-input fold change corresponding to the identified peaks in CC were visualized using CoolBox^84^.

### From regions to fragments

To standardize region lengths for downstream analysis, unevenly sized library regions were converted to uniformly sized 500 bp overlapping fragments with a 50 bp step size, using windowmaker function of BedTools.

### Differential activity analysis

To calculate differential enhancer activity between each KO library and CC, read depth of the 500 bp fragments was calculated from the filtered bam files of each library replicate using coverage utility from BedTools. The generated read depth files for the KO and CC library replicates were then used to identify differentially active fragments using DESeq2 with default parameters^85^. Differentially activity of a fragment is quantified by their DESeq2 assigned log2 Fold Change in KO line compared to CC.

### Defining fragment categories

Fragments were categorized by their activity and response i.e. differential activity as follows: (1) Active and Inactive fragments: Fragments which had >95% overlap with library peaks were termed active for that library while those which do not were termed inactive, (2) Responsive and Unresponsive fragments: Fragments which were significantly differentially active (p-adjusted<0.01) in a KO library compared to CC were termed responsive whereas those which do not exhibit differential activity were termed non-responsive, (3) Induced and Repressed fragments: Responsive fragments which had a positive log2 Fold Change value in the KO line compared to CC were termed induced while those with negative value were termed repressed, (4) Gained and Lost fragments: Induced fragments which were also classified as active in the KO library but inactive in CC were termed gained whereas repressed fragments which were inactive in KO line but active in CC were termed lost, (5) Always active and always inactive fragments: Fragments which were consistently classified as active in all the libraries including CC were termed always active whereas fragments that are consistently inactive in all the libraries including CC were termed always inactive. We visualized the different category of fragments by creating “activity maps” that depict a fragment’s response to a KO compared to its activity in CC. Response of a fragment to a KO is quantified by the DESeq2 calculated fold change of KO reads over CC reads and its activity in CC is quantified by the RPKM normalized fold change of CC reads over input reads in log2 scale. The intersect utility from BedTools was used to determine any overlap between peaks and fragments.

### Sequence determinant analysis

#### (A) Motif enrichment analysis

Motif enrichment analysis was performed using HOMER^86^ and MEME^87^ with the fragments within a specific category as the primary regions of interest and fragments from the contrasting category as the background. For example, to assess enrichment among active fragments in a KO library, active fragments of that library served as the primary and inactive fragments from the same library served as the background.

#### (B) Intra-library classification models

Machine learning models were trained to classify active and inactive fragments for each KO libraries as well as CC. For each library, both linear (logistic regression) and deep learning (Residual Networks) models were trained to classify the fragments based on their library specific activity. For each classification problem, training, validation, and test data were created by splitting the fragments randomly into three parts comprising of 70%, 15% and 15% fragments. Models were implemented in PyTorch and model weights were optimized with Adam to minimize binary cross-entropy loss. We used an initial learning rate of 0.001 and a batch size of 64. Model performance was evaluated on the held-out test data by calculating average precision score, precision, recall, balanced accuracy, F1-score, and area under the receiver operating characteristic curve using scikit-learn.

##### (a) Logistic Regression

For the logistic regression model, fragments were numerically encoded by scanning for presence of known motifs from the HOCOMOCO database (v11 CORE collection) using the findMotifsGenome utility from HOMER. Each fragment was represented by a vector whose length equals the number of TF motifs present in the database and the values denote probability scores of presence of TF binding motifs in that fragment. The logistic regression-based model implemented in PyTorch takes the motif derived features of a fragment as input and predicts activity of that fragment. After training, the motifs were ranked based on the trained model weights assigned to each TF motif to estimate their contribution towards accurate classification of fragment activity. The most relevant motifs for fragment classification of each library were visualized as a heatmap using seaborn \cite.

##### (b) Deep Learning

Deep learning models are capable of automatic feature learning directly from nucleotide sequences. For deep learning models, the fragments were one hot encoded and served as model inputs. The deep learning model architecture was similar to a previously published method^88^ with the input length of the sequence set at 1000 bp. The model comprised two parts: an encoder that learns sequence features from raw DNA and a classifier that converts the encoder output into a probability score denoting fragment activity. The encoder consisted of multiple convolutional layers followed by residual blocks, with batch normalization and max-pooling layers in between. The classifier contained two fully connected layers with ReLU activation and batch normalization. After training, interpretable attribution scores were generated for each test fragment using integrated gradients method implemented in captum within PyTorch. Integrated Gradients assigns an importance score to each nucleotide, indicating its effect on the predicted fragment activity. Thereafter, TF-MoDISco was used to cluster Integrated Gradients derived importance scores into seqlets and aggregate them into consolidated motifs. The resulting motifs were then matched to known TF motifs using TOMTOM^89^, and motif locations were mapped back to fragments to interpret sequence features learned by the model.

### RNA-seq data analysis

Raw RNA-seq FASTQ files were processed with an in-house pipeline. Adapters were trimmed with Trimmomatic^90^, and trimmed reads were aligned to the human reference genome (GRCh38) using the STAR aligner^91^. Gene-level counts were obtained with HTSeq-count^92^. Read quality and replicate concordance were assessed by principal component analysis (PCA) using scikit-learn. All tools were run with default parameters. Differential expression between each knockout (KO) and the control (CC) was tested with DESeq2 using the library-specific count matrices and genes with adjusted p < 0.01 were considered differentially expressed. The percentage of differentially expressed genes per KO was visualized with seaborn.

### Connecting enhancers to target genes

Fragments were linked to putative target genes using (i) a nearest-gene approach and (ii) the Activity-by-Contact (ABC) model^13^. For nearest-gene links, we used BEDTools closest function to assign the nearest annotated gene within 5,000 bp of each fragment. For ABC model, enhancer activity and chromatin contact are the required inputs. The ABC model uses DNase-seq/ATAC-seq and H3K27ac ChIP seq data to calculate a proxy for enhancer activity. Instead of the model’s default activity proxies, we used the normalized output-over-input fold change of STARR-seq reads in the CC library as a measure of native enhancer-activity. ABC scores were computed per fragment-gene pair, and links with ABC score >0.1 were retained as predicted targets.

## ACKNOWLEDGEMENTS

We thank Drs. Istvan Albert, Aswathy Sebastian, Ross Hardison, Craig Praul, and the Penn State Genomics Core Facility for technical support for this project. This work was supported by the National Institutes of Health grants R01-GM121907 (National Institute of General Medical Sciences) and R21-NS122398 (National Institute of Neurological Disorders and Stroke), and resources from the Huck Institutes of the Life Sciences to S.G.

## AUTHOR CONTRIBUTIONS

M.D., D.B, and S.G. designed the study and analyses. M.D., A.H., S.M., J.S. and J.M. performed the experiments. D.B., A.H., and M.J. designed bioinformatics pipelines, and performed all statistical, enrichment, and modeling analysis. S.G., A.G., and H.S. provided reagents and facilities to perform experiments. M.D., D.B., and S.G. interpreted the results and wrote the manuscript with approval from all authors.

## DATA AVAILABILITY

All sequence data generated in this study have been submitted to the NCBI BioProject database (https://www.ncbi.nlm.nih.gov/bioproject/) under accession number PRJNA879724. Processed data and summary statistics generated in this study are provided in **Supplementary Information**.

## CODE AVAILABILITY

All source code for preprocessing and analysis of sequencing data generated in this study is available on GitHub (https://github.com/deeprob/starrseq_results).

## REFERENCES

1. Shlyueva, D., Stampfel, G. & Stark, A. Transcriptional enhancers: from properties to genome-wide predictions. Nat. Rev. Genet. 15, 272–286 (2014).

2. Schoenfelder, S. & Fraser, P. Long-range enhancer-promoter contacts in gene expression control. Nat. Rev. Genet. 20, 437–455 (2019).

3. Lambert, S. A. et al. The Human Transcription Factors. Cell 172, 650–665 (2018).

4. Banerji, J., Olson, L. & Schaffner, W. A lymphocyte-specific cellular enhancer is located downstream of the joining region in immunoglobulin heavy chain genes. Cell 33, 729–740 (1983).

5. Verfaillie, A. et al. Multiplex enhancer-reporter assays uncover unsophisticated TP53 enhancer logic. Genome Res. 26, 882–895 (2016).

6. Liu, S. et al. Systematic identification of regulatory variants associated with cancer risk. Genome Biol. 18, 194 (2017).

7. Shlyueva, D. et al. Hormone-responsive enhancer-activity maps reveal predictive motifs, indirect repression, and targeting of closed chromatin. Mol. Cell 54, 180–192 (2014).

8. Johnson, G. D. et al. Human genome-wide measurement of drug-responsive regulatory activity. Nat. Commun. 9, 5317 (2018).

9. Ma, S. et al. Chromatin Potential Identified by Shared Single-Cell Profiling of RNA and Chromatin. Cell 183, 1103–1116.e20 (2020).

10. Chen, S., Lake, B. B. & Zhang, K. High-throughput sequencing of the transcriptome and chromatin accessibility in the same cell. Nat. Biotechnol. 37, 1452–1457 (2019).

11. Cao, J. et al. Joint profiling of chromatin accessibility and gene expression in thousands of single cells. Science 361, 1380–1385 (2018).

12. Javierre, B. M. et al. Lineage-Specific Genome Architecture Links Enhancers and Non-coding Disease Variants to Target Gene Promoters. Cell 167, 1369–1384.e19 (2016).

13. Fulco, C. P. et al. Activity-by-contact model of enhancer-promoter regulation from thousands of CRISPR perturbations. Nat. Genet. 51, 1664–1669 (2019).

14. Avsec, Ž., et al. Base-resolution models of transcription-factor binding reveal soft motif syntax. Nat. Genet. 53, 354–366 (2021).

15. Gasperini, M. et al. A Genome-wide Framework for Mapping Gene Regulation via Cellular Genetic Screens. Cell 176, 377–390.e19 (2019).

16. Andersson, R. et al. An atlas of active enhancers across human cell types and tissues. Nature 507, 455–461 (2014).

17. Schöne, S. et al. Synthetic STARR-seq reveals how DNA shape and sequence modulate transcriptional output and noise. PLoS Genet. 14, e1007793 (2018).

18. Kalita, C. A. et al. High-throughput characterization of genetic effects on DNA-protein binding and gene transcription. Genome Res. 28, 1701–1708 (2018).

19. Arnold, C. D. et al. Genome-wide quantitative enhancer activity maps identified by STARR-seq. Science 339, 1074–1077 (2013).

20. Ran, F. A. et al. Genome engineering using the CRISPR-Cas9 system. Nat. Protoc. 8, 2281–2308 (2013).

21. Vanhille, L. et al. High-throughput and quantitative assessment of enhancer activity in mammals by CapStarr-seq. Nat. Commun. 6, 6905 (2015).

22. Neumayr, C., Pagani, M., Stark, A. & Arnold, C. D. STARR-seq and UMI-STARR-seq: Assessing Enhancer Activities for Genome-Wide-, High-, and Low-Complexity Candidate Libraries. Curr. Protoc. Mol. Biol. 128, e105 (2019).

23. MacConaill, L. E. et al. Unique, dual-indexed sequencing adapters with UMIs effectively eliminate index cross-talk and significantly improve sensitivity of massively parallel sequencing. BMC Genomics 19, 30 (2018).

24. Das, M., Hossain, A., Banerjee, D., Praul, C. A. & Girirajan, S. Challenges and considerations for reproducibility of STARR-seq assays. Genome Res. 33, 479–495 (2023).

25. Klein, J. C. et al. A systematic evaluation of the design and context dependencies of massively parallel reporter assays. Nat. Methods 17, 1083–1091 (2020).

26. Lee, D. et al. STARRPeaker: uniform processing and accurate identification of STARR-seq active regions. Genome Biol. 21, 298 (2020).

27. Boyle, A. P. et al. High-resolution mapping and characterization of open chromatin across the genome. Cell 132, 311–322 (2008).

28. Verheul, T. C. J., van Hijfte, L., Perenthaler, E. & Barakat, T. S. The Why of YY1: Mechanisms of Transcriptional Regulation by Yin Yang 1. Front. Cell Dev. Biol. 8, 592164 (2020).

29. Chandra, J., Kuo, P. T. Y., Hahn, A. M., Belz, G. T. & Frazer, I. H. Batf3 selectively determines acquisition of CD8+ dendritic cell phenotype and function. Immunol. Cell Biol. 95, 215–223 (2017).

30. Ataide, M. A. et al. BATF3 programs CD8+ T cell memory. Nat. Immunol. 21, 1397–1407 (2020).

31. Beanan, M. J. & Sargent, T. D. Regulation and function of Dlx3 in vertebrate development. Dev. Dyn. Off. Publ. Am. Assoc. Anat. 218, 545–553 (2000).

32. Olbrot, M., Rud, J., Moss, L. G. & Sharma, A. Identification of beta-cell-specific insulin gene transcription factor RIPE3b1 as mammalian MafA. Proc. Natl. Acad. Sci. U. S. A. 99, 6737–6742 (2002).

33. Watson, G., Ronai, Z. & Lau, E. ATF2, a paradigm of the multifaceted regulation of transcription factors in biology and disease. Pharmacol. Res. 119, 347–357 (2017).

34. Chriett, S. et al. SCRT1 is a novel beta cell transcription factor with insulin regulatory properties. Mol. Cell. Endocrinol. 521, 111107 (2021).

35. Lea, A. J. et al. Genome-wide quantification of the effects of DNA methylation on human gene regulation. eLife 7, e37513 (2018).

36. de Almeida, B. P., Reiter, F., Pagani, M. & Stark, A. DeepSTARR predicts enhancer activity from DNA sequence and enables the de novo design of synthetic enhancers. Nat. Genet. 54, 613–624 (2022).

37. Farley, E. K., Olson, K. M., Zhang, W., Rokhsar, D. S. & Levine, M. S. Syntax compensates for poor binding sites to encode tissue specificity of developmental enhancers. Proc. Natl. Acad. Sci. U. S. A. 113, 6508–6513 (2016).

38. Grossman, S. R. et al. Systematic dissection of genomic features determining transcription factor binding and enhancer function. Proc. Natl. Acad. Sci. U. S. A. 114, E1291–E1300 (2017).

39. Smith, G. D., Ching, W. H., Cornejo-Páramo, P. & Wong, E. S. Decoding enhancer complexity with machine learning and high-throughput discovery. Genome Biol. 24, 116 (2023).

40. Grumolato, L. et al. β-Catenin-independent activation of TCF1/LEF1 in human hematopoietic tumor cells through interaction with ATF2 transcription factors. PLoS Genet. 9, e1003603 (2013).

41. Lopez-Bergami, P., Lau, E. & Ronai, Z. Emerging roles of ATF2 and the dynamic AP1 network in cancer. Nat. Rev. Cancer 10, 65–76 (2010).

42. Fang, H. et al. Trans- and cis-acting effects of Firre on epigenetic features of the inactive X chromosome. Nat. Commun. 11, 6053 (2020).

43. Siggens, L., Cordeddu, L., Rönnerblad, M., Lennartsson, A. & Ekwall, K. Transcription-coupled recruitment of human CHD1 and CHD2 influences chromatin accessibility and histone H3 and H3.3 occupancy at active chromatin regions. Epigenetics Chromatin 8, 4 (2015).

44. Ramasamy, S. et al. The Mediator complex regulates enhancer-promoter interactions. Nat. Struct. Mol. Biol. 30, 991–1000 (2023).

45. Fournier, M. et al. FOXA and master transcription factors recruit Mediator and Cohesin to the core transcriptional regulatory circuitry of cancer cells. Sci. Rep. 6, 34962 (2016).

46. Sahu, B. et al. Sequence determinants of human gene regulatory elements. Nat. Genet. 54, 283–294 (2022).

47. He, K., Zhang, X., Ren, S. & Sun, J. Deep Residual Learning for Image Recognition. Preprint at 10.48550/ARXIV.1512.03385 (2015).

48. Shrikumar, A., et al. Technical Note on Transcription Factor Motif Discovery from Importance Scores (TF-MoDISco) version 0.5.6.5. Preprint at 10.48550/ARXIV.1811.00416 (2018).

49. Hai, T. & Curran, T. Cross-family dimerization of transcription factors Fos/Jun and ATF/CREB alters DNA binding specificity. Proc. Natl. Acad. Sci. U. S. A. 88, 3720–3724 (1991).

50. Fawcett, T. W., Martindale, J. L., Guyton, K. Z., Hai, T. & Holbrook, N. J. Complexes containing activating transcription factor (ATF)/cAMP-responsive-element-binding protein (CREB) interact with the CCAAT/enhancer-binding protein (C/EBP)-ATF composite site to regulate Gadd153 expression during the stress response. Biochem. J. 339 **( Pt** **1**), 135–141 (1999).

51. Kerppola, T. K. & Curran, T. Fos-Jun heterodimers and Jun homodimers bend DNA in opposite orientations: implications for transcription factor cooperativity. Cell 66, 317–326 (1991).

52. Pizzo, L. et al. Rare variants in the genetic background modulate cognitive and developmental phenotypes in individuals carrying disease-associated variants. Genet. Med. Off. J. Am. Coll. Med. Genet. 21, 816–825 (2019).

53. Jensen, M. et al. Genetic modifiers and ascertainment drive variable expressivity of complex disorders. Cell S0092-8674(25)01080–3 (2025) doi:10.1016/j.cell.2025.09.012.

54. Muir, A. M. et al. Variants in GNAI1 cause a syndrome associated with variable features including developmental delay, seizures, and hypotonia. Genet. Med. Off. J. Am. Coll. Med. Genet. 23, 881–887 (2021).

55. Huang, Y. et al. Novel dominant and recessive variants in human ROBO1 cause distinct neurodevelopmental defects through different mechanisms. Hum. Mol. Genet. 31, 2751–2765 (2022).

56. Otto, F., Kanegane, H. & Mundlos, S. Mutations in the RUNX2 gene in patients with cleidocranial dysplasia. Hum. Mutat. 19, 209–216 (2002).

57. O’Roak, B. J. et al. Sporadic autism exomes reveal a highly interconnected protein network of de novo mutations. Nature 485, 246–250 (2012).

58. Kim, K. M. et al. A new MIF4G domain-containing protein, CTIF, directs nuclear cap-binding protein CBP80/20-dependent translation. Genes Dev. 23, 2033–2045 (2009).

59. Wang, S. & Sun, S. Translation dysregulation in neurodegenerative diseases: a focus on ALS. Mol. Neurodegener. 18, 58 (2023).

60. Jaffrey, S. R. & Wilkinson, M. F. Nonsense-mediated RNA decay in the brain: emerging modulator of neural development and disease. Nat. Rev. Neurosci. 19, 715–728 (2018).

61. Sun, J. et al. An integrated framework for functional dissection of variable expressivity in genetic disorders. MedRxiv Prepr. Serv. Health Sci. 2025.07.22.25331885 (2025) doi:10.1101/2025.07.22.25331885.

62. Vockley, C. M. et al. Massively parallel quantification of the regulatory effects of noncoding genetic variation in a human cohort. Genome Res. 25, 1206–1214 (2015).

63. Liu, Y. et al. Functional assessment of human enhancer activities using whole-genome STARR-sequencing. Genome Biol. 18, 219 (2017).

64. Tewhey, R. et al. Direct Identification of Hundreds of Expression-Modulating Variants using a Multiplexed Reporter Assay. Cell 165, 1519–1529 (2016).

65. Lee, D. et al. Massively parallel reporter assays identify functional enhancer variants at QT interval GWAS loci. BioRxiv Prepr. Serv. Biol. 2025.03.11.642686 (2025) doi:10.1101/2025.03.11.642686.

66. Ulirsch, J. C. et al. Systematic Functional Dissection of Common Genetic Variation Affecting Red Blood Cell Traits. Cell 165, 1530–1545 (2016).

67. Catizone, A. N. et al. Locally acting transcription factors regulate p53-dependent cis-regulatory element activity. Nucleic Acids Res. 48, 4195–4213 (2020).

68. Huang, D. & Ovcharenko, I. Enhancer-silencer transitions in the human genome. Genome Res. 32, 437–448 (2022).

69. Cui, X. et al. CREATE: cell-type-specific cis-regulatory element identification via discrete embedding. Nat. Commun. 16, 4607 (2025).

70. Zhu, X. et al. Uncovering the whole genome silencers of human cells via Ss-STARR-seq. Nat. Commun. 16, 723 (2025).

71. Siersbæk, R. et al. Transcription factor cooperativity in early adipogenic hotspots and super-enhancers. Cell Rep. 7, 1443–1455 (2014).

72. Pott, S. & Lieb, J. D. What are super-enhancers? Nat. Genet. 47, 8–12 (2015).

73. Su, C., Lee, D., Jin, P. & Zhang, J. scMultiMap: Cell-type-specific mapping of enhancers and target genes from single-cell multimodal data. Nat. Commun. 16, 3941 (2025).

74. Sakaue, S. et al. Tissue-specific enhancer-gene maps from multimodal single-cell data identify causal disease alleles. Nat. Genet. 56, 615–626 (2024).

75. Li, Y. et al. Enhancer-driven gene regulatory networks inference from single-cell RNA-seq and ATAC-seq data. Brief. Bioinform. 25, bbae369 (2024).

76. Motallebnejad, P., et al. Integrated ATAC-seq and RNA-seq analysis identifies key regulatory elements in NK cells activated with feeder cells and IL-2. Bioeng. Transl. Med. 10, e10747 (2025).

77. Bai, Y. et al. Integrative analysis based on ATAC-seq and RNA-seq reveals a novel oncogene PRPF3 in hepatocellular carcinoma. Clin. Epigenetics 16, 154 (2024).

78. Inoue, F. et al. A systematic comparison reveals substantial differences in chromosomal versus episomal encoding of enhancer activity. Genome Res. 27, 38–52 (2017).

79. Muerdter, F. et al. Resolving systematic errors in widely used enhancer activity assays in human cells. Nat. Methods 15, 141–149 (2018).

80. Quinlan, A. R. & Hall, I. M. BEDTools: a flexible suite of utilities for comparing genomic features. Bioinforma. Oxf. Engl. 26, 841–842 (2010).

81. Li, H. & Durbin, R. Fast and accurate short read alignment with Burrows-Wheeler transform. Bioinforma. Oxf. Engl. 25, 1754–1760 (2009).

82. Li, H. et al. The Sequence Alignment/Map format and SAMtools. Bioinforma. Oxf. Engl. 25, 2078–2079 (2009).

83. Ramírez, F., Dündar, F., Diehl, S., Grüning, B. A. & Manke, T. deepTools: a flexible platform for exploring deep-sequencing data. Nucleic Acids Res. 42, W187–191 (2014).

84. Xu, W. et al. CoolBox: a flexible toolkit for visual analysis of genomics data. BMC Bioinformatics 22, 489 (2021).

85. Love, M. I., Huber, W. & Anders, S. Moderated estimation of fold change and dispersion for RNA-seq data with DESeq2. Genome Biol. 15, 550 (2014).

86. Heinz, S. et al. Simple combinations of lineage-determining transcription factors prime cis-regulatory elements required for macrophage and B cell identities. Mol. Cell 38, 576–589 (2010).

87. Bailey, T. L. et al. MEME SUITE: tools for motif discovery and searching. Nucleic Acids Res. 37, W202–208 (2009).

88. Nair, S., Kim, D. S., Perricone, J. & Kundaje, A. Integrating regulatory DNA sequence and gene expression to predict genome-wide chromatin accessibility across cellular contexts. Bioinforma. Oxf. Engl. 35, i108–i116 (2019).

89. Gupta, S., Stamatoyannopoulos, J. A., Bailey, T. L. & Noble, W. S. Quantifying similarity between motifs. Genome Biol. 8, R24 (2007).

90. Bolger, A. M., Lohse, M. & Usadel, B. Trimmomatic: A flexible trimmer for Illumina sequence data. Bioinformatics 30, 2114–2120 (2014).

91. Dobin, A. et al. STAR: ultrafast universal RNA-seq aligner. Bioinforma. Oxf. Engl. 29, 15–21 (2013).

92. Anders, S., Pyl, P. T. & Huber, W. HTSeq--a Python framework to work with high-throughput sequencing data. Bioinforma. Oxf. Engl. 31, 166–169 (2015).

