## Supplemental File for "Quantifying the impact of genetic mutations on enhancer dynamics"

### Supplementary Figures

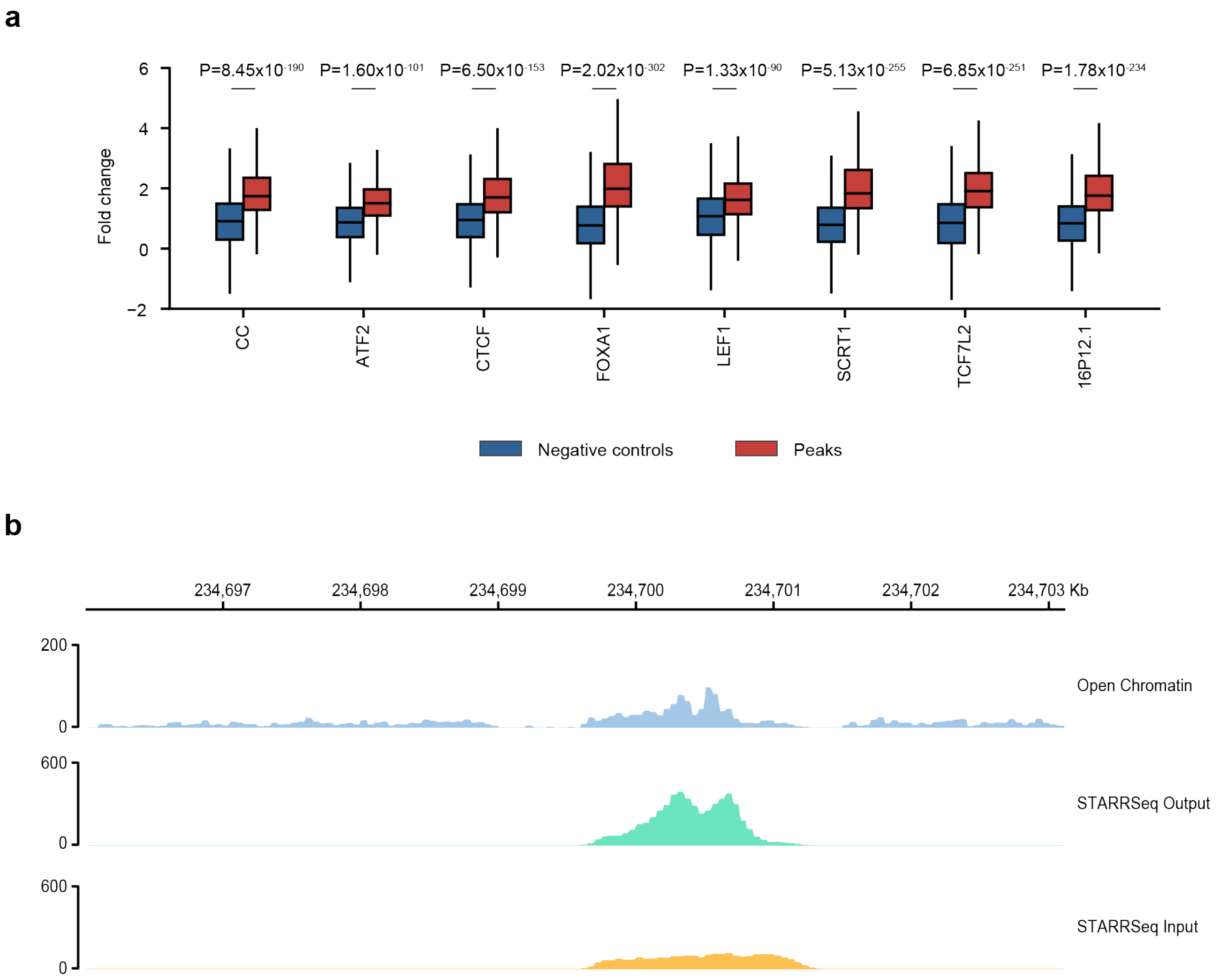

**Supplementary Figure 1: Quality control of STARR-seq peaks**

**(a)** Box plot illustrating STARR-seq readout (RPKM normalized Output/Input reads) for negative controls (fragments overlapping exonic regions) and STARRPeaker called peaks across KO libraries. **(b)** Demonstrating read pileup at a STARRPeaker called peak measured through STARR-seq assay that overlapped an open chromatin region measured through DNase-seq.

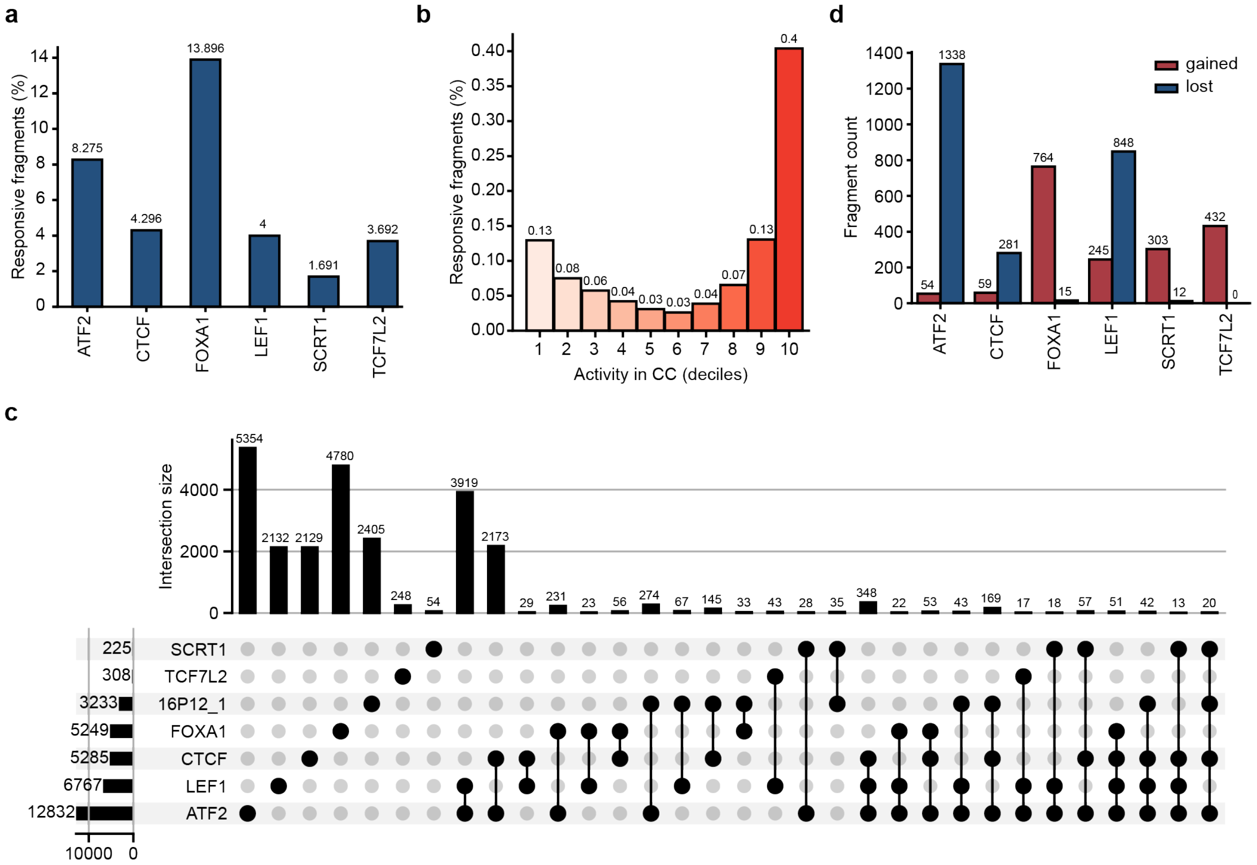

**Supplementary Figure 2: Responsive fragments across TF-KO libraries**

**(a)** Bar plot showing the percentage of responsive fragments across KO libraries. **(b)** Bar plot showing the percentage of responsive fragments stratified by CC activity in the ATF2 KO line. **(c)** Upset plot showing the number repressed fragments that overlapped between all KO lines. **(d)** Bar plot showing the number of gained and lost fragments across KO lines.

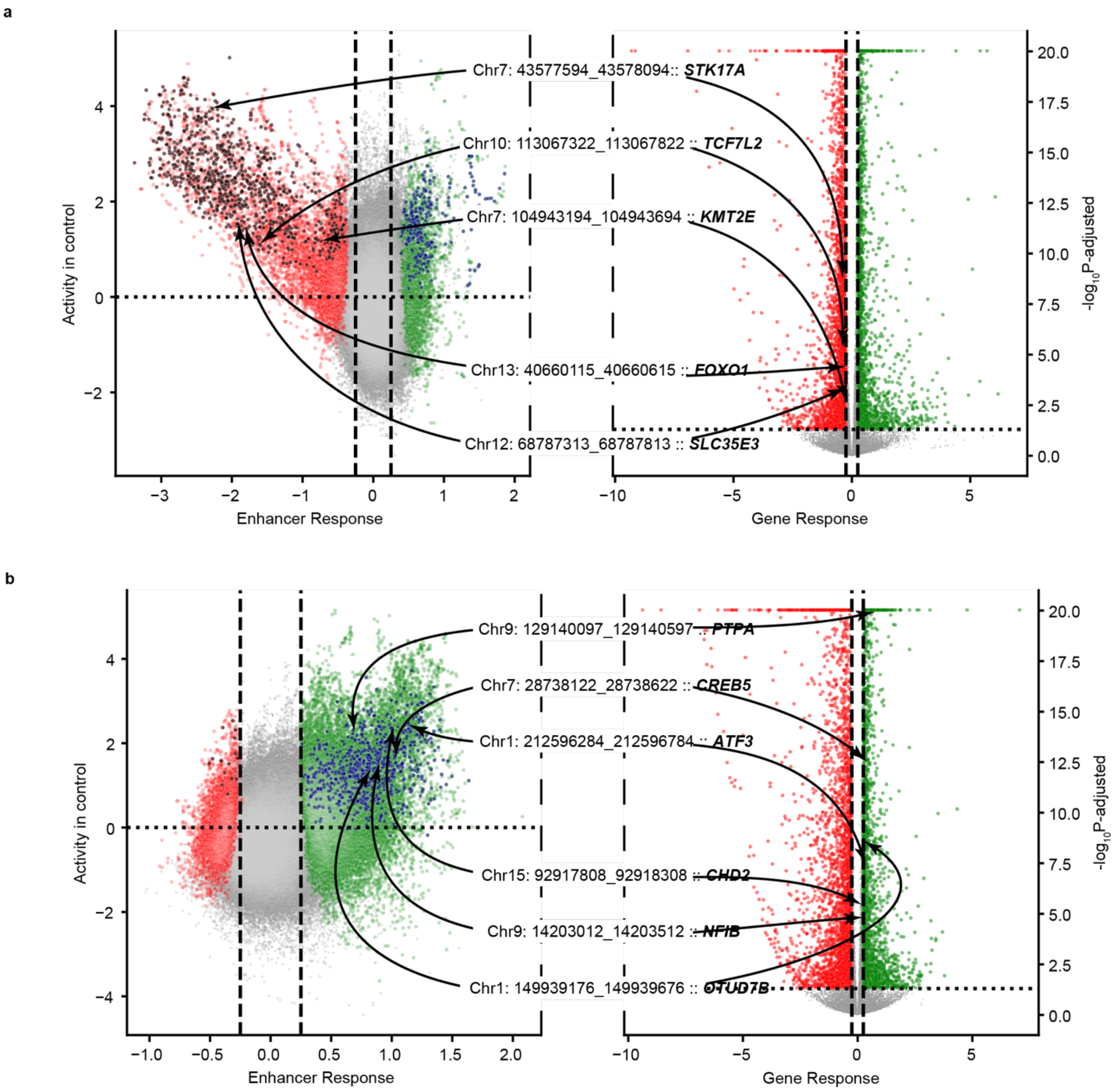

**Supplementary Figure 3: Mapping enhancers to target genes**

Differential enhancer activity map measuring log2 fold change of normalized output reads between KO and CC compared to CC activity of fragments and corresponding volcano plot illustrating differentially expressed target genes in **(a)** LEF1 and **(b)** FOXA1 KO lines.

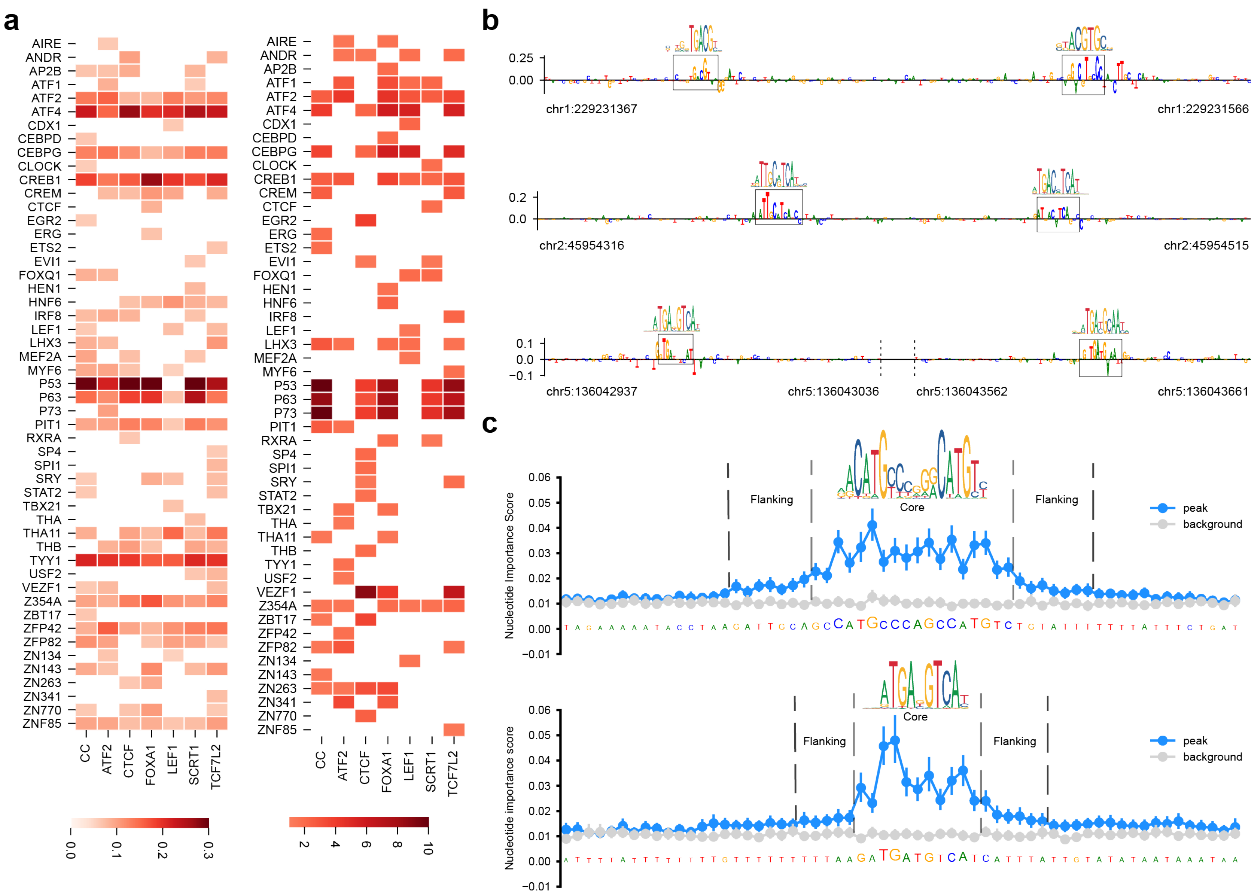

**Supplementary Figure 4: Learning the regulatory code**

**(a)** Heatmap illustrating the importance score of common motifs identified by the linear (left) and deep learning (right) models to predict enhancer activity across TF KO lines. **(b)** Combinations of motifs identified by the deep learning model to accurately predict active enhancers which the linear model missed. **(c)** Importance score of flanking sequence highlighted by the deep learning model to accurately predict enhancer activity of peaks which were missed by the linear model.

**Supplementary Data inventory**

**Supplementary Data 1**: Activity, STARRPeaker peak status in CC and all KO lines, differential activity statistics in KO lines compared to CC and defined category of fragments with respect to CC and KO lines.

**Supplementary Data 2**: Motif Enrichment Analysis results for all categories of fragments across CC and KO lines.

**Supplementary Data 3**: Differentially active enhancers mapped to their differentially expressed target genes along with differential expression statistics across KO lines.

**Transcription Factors selected in this study**

**Supplementary Table 1**: Shortlisted enhancer-binding transcription factors and their properties

| Transcription Factor | Total ChIP-seq sites (HEK293) | Library Overlapping sites | TF Function | References |
|---|---|---|---|---|
| FOXA1 | 1,272 | 595 | • Pioneer TF responsible for opening condensed chromatin to enable further TF binding and gene activation.<br>• Associated with nuclear receptors in specific cell-types such as ESR1 in MCF-7 cells and AR in LNCaP cells. | [1]<br>[2]<br>[3] |
| ATF2 | 30,810 | 5,727 | • TF is a member of the AP1 family of regulators and activates gene expression through homo/heterodimerization with other AP1 members such as Jun, Fos or CREB.<br>• It is involved in cell physiology, development, signaling and is associated with tumorigenesis. | [4]<br>[5]<br>[6] |
| CTCF | 51,527 | 2,591 | • Known insulator binding protein found enriched at TAD boundaries.<br>• CTCF binding sites are also found across enhancers within TADs. | [7]<br>[8] |
| LEF1 | 3,683 | 5,98 | • Forms transcriptional complex through binding of T-cell receptor alpha enhancer.<br>• Interacts with nuclear CTNNB1 and activates wnt signaling genes associated with cell cycle, survival and tumorigenesis. | [9]<br>[10] |
| TCF7L2 | 8,961 | 492 | • Involved in CTNNB1 mediated transcriptional activation or repression. Mediates downstream effects of wnt signaling. | [11]<br>[12] |

| | | | | |
|---|---|---|---|---|
| | | | • Has cell-type specific co-localization properties affecting gene regulation with factors like GATA3 in MCF7 and HNF4α and FOXA2 in HepG2 cell lines. | |
| SCRT1 | 26,948 | 5,310 | • Known transcriptional repressor that controls glucose induced insulin secretion in β-cells.<br>• It is further repressed by REST, an inhibitor that is associated with neurogenesis and neuronal differentiation. | [13] [14] [15] |

**Generation of CRISPR/Cas9 mediated TF-depletion lines**

We selected HEK293T cells as the model organism of choice due to ease of culture and due to expression of a wide range of TFs. We used a dual sgRNA strategy (Tai et al 2016) for engineering the deletions. Each sgRNA pair was designed to target regions immediately upstream or downstream of the TSS and regions immediately upstream, downstream or within the terminal exon with the intention of cutting the entire genomic region in between for a complete deletion.

<u>*Single guide RNA design*</u>

We designed all sgRNA using built-in tools on Benchling (San Francisco, CA). We selected guides based on off-target and on-target scores (from Benchling) as well as on sequence similarities (or lack thereof) with the human genome, determined through BLAT (Kent 2002) (The BLAST Like Alignment Tool) available on UCSC genome browser. All sgRNA used are provided in **Supplementary Table 2**. Guides were designed to include overhang sequences to facilitate sgRNA cloning into CRISPR/Cas9 vectors according to protocols for PX330 backbone containing plasmids provided by the Zhang lab (Ran et al 2013).

**Supplementary Table 2**: List of sgRNA

| Transcription Factor | gRNA (5'-3') |
|---|---|
| ATF2_1 | `TTACTGTTACTATATGGGGA` |
| ATF2_2 | `GATGCAAAGTGAGTGCAGAG` |

| | |
|---|---|
| FOXA1_1 | TTGTGGGATAACTGACCCCG |
| FOXA1_2 | AGGATGGAGTTCATACACAA |
| CTCF_1 | CTTTTGAAAGTTGGCGCCCG |
| CTCF_2 | CTAGAGAAAGTACCATCCCA |
| SCRT1_1 | CGCCGGCAGCTTCAGCACCG |
| SCRT1_2 | GGGTGGGAATATGTACAAGG |
| LEF1_1 | TAAAACGGACATCTCCAGCG |
| LEF1_2 | AGGAATTGGAAAGGTTCCAG |
| TCF7L2_1 | TCAACTCACTCAAATCCGAG |
| TCF7L2_2 | AAACCATAAACAAAGCAGCG |
| 16p12.1_1 | TCGGTGCTTAGGATCAGCCT |
| 16p12.1_2 | CAACCATGTCAGCTAGTGGC |

*Vector preparation and sgRNA cloning*

We selected two separate vectors for delivering the two guides. Vector A, pSpCas9(BB)-2A-Puro (PX459) V2.0 was a gift from Feng Zhang (Addgene plasmid # 62988; http://n2t.net/addgene:62988; RRID: Addgene_62988) (Ran et al 2013). This vector contained Cas9, an sgRNA scaffold, a puromycin resistance marker and an ampicillin resistance marker. Vector B, pGH020_sgRNA_G418-GFP was a gift from Michael Bassik (Addgene plasmid # 85405; http://n2t.net/addgene:85405; RRID: Addgene_85405) (Hess et al 2016). This vector contained cloning sites for sgRNA, a G418 resistance marker, an ampicillin marker and an EGFP sequence for expressing green fluorescent protein. However, this vector did not contain an sgRNA scaffold sequence which had to be synthesized along with the designed guide RNA sequence for guides to be cloned into this vector **(Supplementary Table 2)**. The vector stocks (addgene, Watertown, MA) were streaked onto LB agar plates with 100 µg/ml ampicillin to isolate colonies and grown overnight. Multiple colonies were used to inoculate 50 ml LB broth with 100 µg/ml ampicillin and grown over night on a shaker incubator at 37ºC and 250 rpm. Plasmid DNA was isolated using ZymoPURE II Plasmid Midiprep kit (Catalog #D4200, Zymo Research, CA). For cloning in guides, vectors were digested with BbsI-HF (Catalog #R3539S, New England Biolabs, MA) restriction enzyme according to manufacturer's protocol. Digested vectors were purified using DNA Clean & Concentrator kit (Catalog #D4033, Zymo Research, CA).

All selected sgRNA were synthesized from IDT (Integrated DNA Technologies, Coralville, IA) as single stranded 25 nmol oligos with standard desalting. Single stranded oligo pairs were annealed according to the Zhang lab protocol, https://www.addgene.org/crispr/zhang/ (Ran et al 2013). Vectors and guides were ligated using Quick Ligase (Catalog #M2200S, New England Biolabs, MA) according to manufacturer's protocol and transformed in NEB 5-alpha Competent *E. coli* (Catalog #C2988J, New England Biolabs, MA) and plated on LB agar plates with 100 µg/ml ampicillin and grown overnight. Multiple transformants were further grown overnight in 4 ml LB broth with 100 µg/ml ampicillin at 37ºC and 250 rpm on a shaker incubator. Plasmid DNA was isolated using GeneJET Plasmid Miniprep Kit (Catalog #K0502, ThermoFisher Scientific, USA) and sequenced using sanger sequencing at the Genomics Core Facility, Huck Institutes of Life Sciences, Pennsylvania State University to validate cloning.

*Transfection HEK293T cells and CRISPR/Cas9 KO validation*

HEK293T cells were cultured according to ATCC guidelines on 6-welled plates up to two passages. We transfected individual wells (Day 1) with both vectors for each TF deletion simultaneously at ~60% confluency using Lipofectamine 3000 following manufacturer's protocol and incubated the cells in OptiMEM for 24 hours. A control line was generated by transfecting empty (no gRNA) vectors called CRISPR Control (CC). We replaced OptiMEM with complete DMEM media (Day 2) and incubated cells for another 24 hours, until cells reached >95% confluency. We validated transfection by visualizing GFP expression and then replaced media with complete DMEM supplemented with Puromycin (1µg/µl) and G418 (700µg/µl) antibiotic. Cells were passaged on day 4 into fresh 6-welled plates and cultured for 10 days. Antibiotic supplemented media was changed and replenished every 3 days. At day 14, media was again replaced by complete DMEM (without antibiotics) for cell recovery. Stocks of each transfected line was frozen upon reaching confluency. Cells in culture were used to isolate genomic DNA (catalog #N170, Sigma Aldrich, MO, USA).

*CRISPR/Cas9 KO assessment and clonal selection*

100 ng of genomic DNA was amplified using genotyping primers designed upstream of left flank gRNA and downstream of right flank gRNA. Successful deletions showed amplified fragment while CC showed no product. Amplified product was sequenced and confirmed using sanger sequencing (data available upon request). Following initial KO confirmation, each KO line was re-cultured and diluted to 50 cells/10ml with 100µl of cells plated per well on a 96 well plate such that each well consisted of a single cell or no cell and grown for 14 days. Single clonal colonies were expanded, and individual clones were re-genotyped and KO clones were re-validated using sanger sequencing.

*Western Blot*

For each TF, all successfully deleted clones were assessed for TF protein depletion using western blot. In brief, cells cultured in 6-welled plates were lysed in RIPA lysis buffer and sonicated. The initial total protein content was estimated using BCA (Bicinchoninic acid) assay (catalog #23225, Thermo Fisher Scientific, MA, USA). 20 µg of total cell lysates were resolved on 10% SDS-PAGE gels and transferred onto 0.2 µm nitrocellulose membranes via Trans-Blot Turbo (BioRad, CA, USA). All blots were then incubated with 5% (w/v) non-fat dry milk in 1X PBS prior to protein detection using TF-specific monoclonal antibodies. GAPDH was used as a background loading control. Protein depletions were compared to protein level on CC line. Clone with maximum TF depletion was selected for STARR-seq and RNA-seq experiments.

**Library transfection quality control**

For generating the output and input screening libraries, we used a slightly scaled and modified version of the UMI STARR Seq protocol (Neumayr et al 2019). We expanded HEK293T cultures onto 24 x 15cm dish (1 dish per replicate) up to 80-90% confluency (~35 x $10^7$ cells). We transfected similarly conditioned replicates on separate days to account for batch effects. Prior to transfection, we aspirated media from each culture dish, washed cells with PBS and OptiMEM reagent (Thermo Fisher # 31985062), added 25 ml of OptiMEM and incubated them at 37ºC for

1 hour. We transfected 38 μg of the STARR-seq plasmid library using 57 μl of Lipofectamine 3000 and 76μl of P3000 enhancer reagent, diluted in 1.5 ml of OptiMEM according to the manufacturers protocol and scaling guidelines (Thermo Fisher #L3000008).

#### References

1. Sérandour, A. A. *et al.* Epigenetic switch involved in activation of pioneer factor FOXA1-dependent enhancers. *Genome Res.* **21**, 555–565 (2011).
2. Jägle, S. *et al. SNAIL1-Mediated Downregulation of FOXA Proteins Facilitates the Inactivation of Transcriptional Enhancer Elements at Key Epithelial Genes in Colorectal Cancer Cells*. *PLoS Genetics* vol. 13 (2017).
3. Friedman, J. R. & Kaestner, K. H. The Foxa family of transcription factors in development and metabolism. *Cell. Mol. Life Sci.* **63**, 2317–2328 (2006).
4. Watson, G., Ronai, Z. & Lau, E. ATF2, a paradigm of the multifaceted regulation of transcription factors in biology and disease. *Pharmacol. Res.* **119**, 347–357 (2017).
5. Huebner, K., Procházka, J., Monteiro, A. C., Mahadevan, V. & Schneider-Stock, R. The activating transcription factor 2: An influencer of cancer progression. *Mutagenesis* **34**, 375–389 (2019).
6. Brockmann, D., Pützer, B. M., Lipinski, K. S., Schmücker, U. & Esche, H. A multiprotein complex consisting of the cellular coactivator p300, AP- 1/ATF, as well as NF-κB is responsible for the activation of the mouse major histocompatibility class I (H-2Kb) enhancer A. *Gene Expr.* **8**, 1–18 (1999).
7. Ren, G. *et al.* CTCF-Mediated Enhancer-Promoter Interaction Is a Critical Regulator of Cell-to-Cell Variation of Gene Expression. *Mol. Cell* **67**, 1049-1058.e6 (2017).
8. Dixon, J. R. *et al.* Topological domains in mammalian genomes identified by analysis of chromatin interactions. *Nature* **485**, 376–380 (2012).
9. Aoki, M., Hecht, A., Kruse, U., Kemler, R. & Vogt, P. K. Nuclear endpoint of Wnt signaling: Neoplastic transformation induced by transactivating lymphoid-enhancing factor 1. *Proc. Natl. Acad. Sci. U. S. A.* **96**, 139–144 (1999).

10. Singhi, A. D. *et al.* Overexpression of Lymphoid Enhancer-Binding Factor 1 (LEF1) in solid-pseudopapillary neoplasms of the pancreas. *Mod. Pathol.* **27**, 1355–1363 (2014).
11. Guo, Q. *et al.* A β-catenin-driven switch in tcf/lef transcription factor binding to dna target sites promotes commitment of mammalian nephron progenitor cells. *eLife* **10**, 1–47 (2021).
12. Frietze, S. *et al.* Cell type-specific binding patterns reveal that TCF7L2 can be tethered to the genome by association with GATA3. *Genome Biol.* **13**, (2012).
13. Chriett, S. *et al.* SCRT1 is a novel beta cell transcription factor with insulin regulatory properties. *Mol. Cell. Endocrinol.* **521**, (2021).
14. Sobel, J. *et al.* Scrt1, a transcriptional regulator of β-cell proliferation identified by differential chromatin accessibility during islet maturation. *Sci. Rep.* **11**, 1–16 (2021).
15. Mandel, G. *et al.* Repressor element 1 silencing transcription factor (REST) controls radial migration and temporal neuronal specification during neocortical development. *Proc. Natl. Acad. Sci. U. S. A.* **108**, 16789–16794 (2011).
